# Digital news consumption as an information foraging problem

**DOI:** 10.64898/2026.11.23.768652

**Authors:** Henrique M. Borges, Ahmed El Hady

## Abstract

News feeds are digitally available all around us, yet the cognitive mechanisms by which we decide what to spend time on, and what to skip, are not well understood. We formulate digital news consumption as an information foraging problem. In this formulation, users move between news articles, formalized as information patches, exploited them like agents consuming food in a canonical patch foraging task. The switching time between news article can be seen as a navigational process that happens at a time akin to the travel time in the canonical patch foraging formalism. To unravel mechanisms determining news reading times, we fit an evidence-accumulation model separately to each of 2,820 readers of a Norwegian newspaper across 1.9 million reads. Reading times are described by accumulation from a variable starting point, in which readers arrive already partly informed. We also show that readers can be divided into two groups reflecting different behavioral patterns that we name: committed and conditional reflecting different distributions of the fitted evidence accumulation model parameters. Our study opens up the opportunity to study the detailed cognitive mechanisms underlying news consumption and our preliminary analysis show that our information foraging approach can be generalized to our types of digital media human interact with on a daily basis.

## 1 Introduction

Modern humans are constantly exposed to digital information in the form of news, posts, videos, etc. The ubiquity of the digital medium around us has potentially shaped how we cognitively process this barrage of information Meshi et al. [2015], Lara and Bokoch [2021]. One pertinent question is how a reader selects which articles to open or interact with while scrolling a feed, and how long to stay with each. Extensive surveys of news habits and sources describe what people consume Barthel et al. [2020], but not the potential cognitive mechanisms that generate these choices. News consumption can be regarded as analogous to resource consumption, with information as the resource, and the move from one article to the next as foraging between patches. Decisions of this type have been studied extensively in behavioural ecology within the framework of optimal foraging theory.

Optimal foraging theory describes how animals allocate time across resource patches by balancing the diminishing rate of return within a patch against the expected gain from travelling to a new one Charnov [1976], Stephens and Krebs [1986]. In order to tackle the potential cognitive mechanism underlying this behavior, recent work by Davidson and El Hady Davidson and El Hady [2019] formalised this trade-off as a *foraging drift-diffusion model* (FDDM), in which patch-leaving decisions arise from a noisy evidence-accumulation process. The FDDM provides a mechanistic account of patch-leaving decisions by linking Charnov’s marginal-value theorem Charnov [1976] with sequential-sampling models from perceptual decision making Ratcliff and McKoon [2008]. In the idealised noiseless limit, Davidson and El Hady derive analytical conditions under which the drift rate and decision threshold reproduce MVT-optimal mean residence times. Under realistic noise conditions, numerical simulations show that the framework additionally generates full residence time distributions whose shape depends on the foraging strategy employed.

Information foraging theory has been proposed as a generalisation of optimal foraging theory, in which the resources consumed are informational rather than nutritional. It was first introduced by Pirolli and Card in a series of seminal papers that established the information scent that guides search, the patch structure of information environments, and the cost-benefit accounting that governs how long a forager stays in an information patch Pirolli and Card [1999], Markant et al. [2013], Pirolli and Card [1995, 1997], Pirolli [1998], Pirolli and Card [1998], Pirolli and Fu [2003], Pirolli [2007, 2003, 2006], Card et al. [1996].

Subsequent work has tested the framework quantitatively in web search, image retrieval and exploratory mobile browsing Liu et al. [2010], Ong [2017], Jaiswal et al. [2023, 2020], Lambros [2005], where it accounts for observed patterns of interaction. The advantage of formulating a generative model is the ability to fit a model to each person performing a behavior, adding quantities that can be used to uncover latent cognitive variables. Similar models have been used to study how saccadic eye movements can be mapped onto a foraging problem Nahari and El Hady [2025] and how agents search for a mate Bernstein and Hady [2026]. The drift-diffusion model, used to formulate those foraging problem, has been used extensively in studying decision making processes Brunton et al. [2013], Piet et al. [2018], Gold and Shadlen [2007]. As mentioned above it has been extended to provide a mechanistic account of foraging behaviours for both individual Davidson and El Hady [2019], Kilpatrick et al. [2021] and social agents Blum Moyse and El Hady [a,b]. The information foraging framework has in turn been used to study perceptual decisions during active visual sampling Ludwig and Evens [2017], which makes a drift-diffusion account a natural choice here. In the majority of studies, evidence-accumulation models have been developed and tested on animals in controlled environments and on humans in laboratory tasks, where the stimuli are designed a priori and the alternatives are known. Whether a leaving rule estimated this way describes decisions people make unprompted, among alternatives nobody arranged, is largely untested and remain an active area of inquiry.

Digital reading records offer a way to test this, as they provide the quantity a first-passage account requires (the time a reader spends in a patch) for enough information patches per person that a model can be fitted to each reader separately rather than to the population. That matters because an individual fit is what allows a parameter to be treated as a property of a person and tested against others in the population, where a fit to the aggregate yields a description of the sample and cannot reveal individual variability in behavioral parameters.

Here we use reading records from a regional Norwegian newspaper to ask whether the decision to leave an article has the structure of an accumulation to a threshold process. We show that reading times can be described by accumulation from a variable starting point, in which readers arrive at an article already partly informed, and this holds in a reader by reader case rather than only in aggregate. We also tested whether the same type of processes (first passage models) can account for the switching time between articles. We found that a constant-start first-passage model does not describe the interval between articles, which is barely distinguishable from a memoryless process, so, the model accounts for certain aspects of foraging behaviour, but does not provide a comprehensive explanation of the entire process. Moreover, the fitted parameters predict behaviour the model was never fitted to, each predicting a different behaviour: a reader’s evidence threshold predicts both how readily they abandon an article and the route they take to the next, while the minimum evidence they gather predicts abandonment alone. The clustering analysis identifies two distinct groups of readers: committed readers, whose initial evidence states are drawn from a restricted range well below the decision threshold, and conditional readers, whose initial evidence states can span the full range from zero to the decision threshold. The dissociation is what justifies claiming the parameters are properties of readers rather than quantities that improve a fit. Our study establish a mechanistic framework to uncover cognitive mechanisms underlying digital news consumption, that can be generalized to other digital platforms humans interact with.

## 2 Results

### 2.1 News consumption is an information foraging process modeled as a bounded evidence accumulation

A reader arriving at a feed is confronted with more articles than they could possibly read and must make two decisions, namely which article to open and when to leave once they have done so (Fig. 1a). In practice, the observed behaviour reduces to a set of durations, but what we seek to explain are the underlying decision mechanisms that produce them. Information foraging theory provides the interpretive framework which connects the two. Framed as a foraging problem, the decision sequence comprises a repeating cycle, where the reader arrives at an information patch (article), exploits it for a given time, and pays a temporal cost in transitioning to the next one. More concretely, and in foraging terms, an article is a patch, the time spent in it is the patch residence time, and the interval between leaving one article and opening the next is the travel time *τ* (Fig. 1b). In this context, the decision to leave is the behaviour to be explained and the distribution of residence times is the behavioral observable.

**Figure 1:**
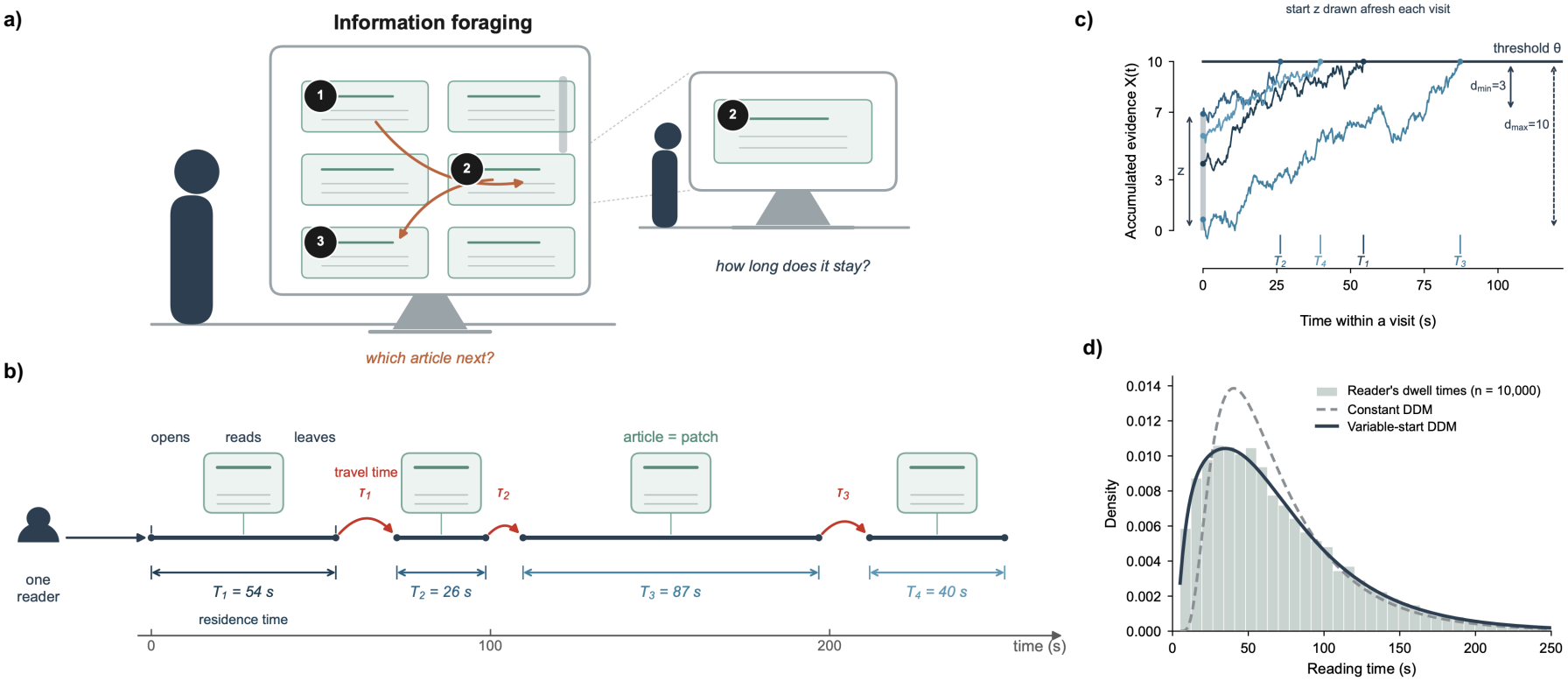
Information Foraging, modelled as bounded evidence accumulation. **(a)** Schematic of the path an individual could take throughout an imaginary session in a news feed, in which the reader makes the decision to open an article (numbered in visit order), whose visit is shown enlarged. **(b)** The same reading session as a patch sequence, an article is a patch, the residence time *T*_*i*_ is the dwell within it, and the travel time *τ*_*i*_ is the interval to the next article. **(c) The mechanism.** Evidence *X*(*t*) accumulates from a starting point *X*(0) = *z*, drawn afresh on each visit from *U* (0, *z*_frac_*θ*), until it reaches the threshold *θ*, at which point the reader leaves. The first-passage times are the residence times in **b**, and the evidence remaining to be gathered lies between *d*_min_ = *θ*(1 − *z*_frac_) and *d*_max_ = *θ*. Sample paths are drawn with the diffusion coefficient, *σ* reduced relative to the fitted regime so that individual trajectories are legible. **(d)** Predicted reading-time densities under two different models, constant-start and variable-start accumulator, with *θ* and *α* held equal so that the effect of start variability is isolated (illustrative parameters, no data are shown).

We model the decision to leave as an evidence-accumulation process (Fig. 1c). Within each visit, a decision variable *X*(*t*) accumulates noisy evidence about the article’s value at rate *α* until it reaches a given threshold *θ*, at which point the reader decides to leave (Methods, Eq. (2)).

Moreover, accumulation does not begin from a state of zero evidence. Indeed, before opening an article, readers have already seen the headline, the associated image, and potentially its source, meaning that some evidence is already available at the outset. Therefore, on each visit, we sample the starting point afresh, leaving a remaining distance to the threshold that varies from visit to visit between *d*_min_ and *θ* (Fig. 1c).

Each reader can then be characterised by three parameters, each with a direct behavioural interpretation. *θ* is the maximum patience, *α* is the rate at which an article produces evidence to that reader and *d*_min_ is the least evidence the reader ever gathers before judging an article, however strongly they arrived disposed to leave it (the assessment floor).

The natural alternative is a constant-start drift-diffusion model, which preserves the accumulation mechanism and the leaving threshold *θ*, and differs only in doing away with the assumption that readers arrive at articles already partly informed. Although similar, the two models predict distinct shapes for the reading-time distribution (Fig. 1d). Under a constant-start, every visit faces the same evidence requirement, so short visits arise so short visits occur only when noise favours an early decision, resulting in a unimodal distribution in which density first increases toward a late mode and then decreases. On the other hand, under a variable-start, the larger *z*_frac_, the more visits begin close to threshold, so the mode moves earlier and the tail thins, while at *z*_frac_ →0 the model reduces to the constant-start case. The two curves in Fig. 1d share *θ* and *α* and differ exclusively in whether the effective distance *d* is fixed or variable, so the difference can be attributed to the starting point alone. These signatures are qualitatively distinct, in particular where they place the mode and how heavily they weight the tail, so reading-time distributions can discriminate between these two models.

### 2.2 Accumulation from a variable starting point predicts News reading times

Reading times in a digital context, and particularly on news websites, are strongly right-skewed, with most article visits ending within a minute and a minority continuing for several. A similar pattern can emerge across multiple families of probability distributions, and a sufficiently flexible distribution will reproduce it without describing any process at all. Therefore, distinguishing the two models of Fig. 1d) is not the same as confirming the existence of a potential evidence accumulation process, therefore we treat the two questions separately. First, we ask whether the starting point varies, which Fig. 2 addresses in three complementary views, and then how an evidence accumulation perspective performs in comparison with descriptions that lack generative content.

**Figure 2:**
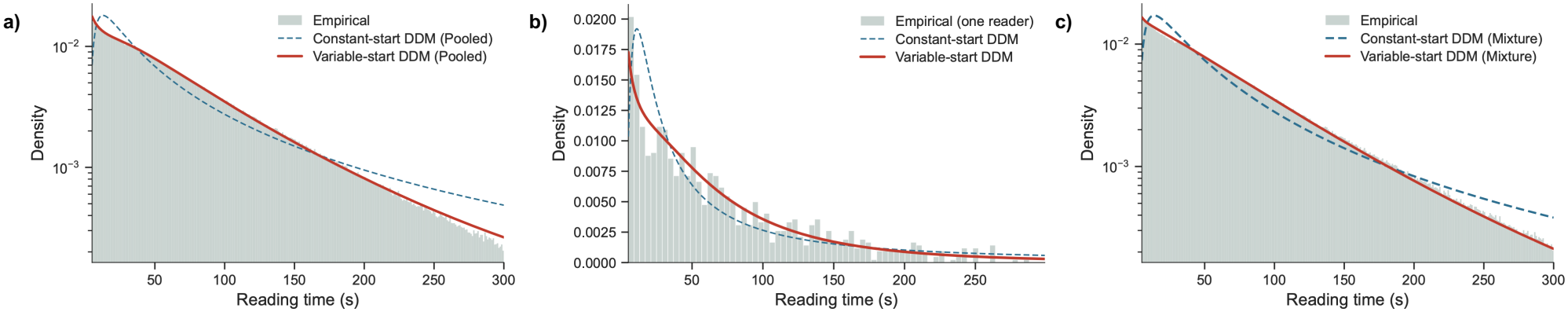
Reading times are described by evidence accumulation from a variable starting point. Two one-sided drift-diffusion models (DDMs) of the decision to leave an article are compared throughout, in which evidence accumulates at a given rate *α* to a threshold *θ*, differing only in the starting point fixed at 0 on every visit (constant-start), or drawn afresh each visit from *U*(0, *z*_frac_*θ*) (variable-start). The diffusion coefficient is fixed at unity, so *θ* and *α* are in units of noise amplitude. **(a) The pooled view**. Reading times pooled across all readers (green, *n* = 1867477 reads), with each model fitted to the pooled sample as though it had come from a single reader (dashed blue, constantstart, and solid red, variable-start), with the densities being conditioned on the [5, 300) s interval. **(b) The individual view**. Reading times of a single reader (green, *n* = 1056 reads), with each model fitted to that reader’s own reads. Selection was done on sample size rather than on goodness of fit, with the reader being selected at the median of fit among readers in the top decile by read count. **(c) The mixture view**. The same pooled reading times as in **(a)** (green), with the read-weighted average of the individually fitted densities, *p*(*t*) = ∑_*i*_ *w*_*i*_*f*_*i*_(*t*), where *w*_*i*_ is proportional to the number of reads contributed by reader *i*. Every parameter derives from a separate per-reader fit, so neither curve in **(c)** has a free parameter at the population level. **(c)** is therefore a prediction, whereas the curves in **(a)** were fitted to the distribution against which they are plotted. Vertical axes are logarithmic in **(a)** and **(c)** and linear in **(b)**. Histogram bars are empirical densities and curves are deterministic given the fitted parameters and carry no sampling error, so no error bars are shown.

To investigate these questions, we started by analyzing all 1867477 reads as if they had come from a single reader. The variable-start model fit attains -5.1280 nats per read against -5.1630 for the constant-start model, and the shapes differ in the way each of the mechanisms predicts (Fig. 2a). One can observe that the direction of the constant-start model’s failure is itself informative. The constantstart density overestimates the frequency of mid-range visits while underestimating the frequency of short visits and a subset of long visits. This pattern is consistent with a first-passage density whose mode occurs later than that of the observed distribution.

A similar contrast can be observed within single readers (Fig. 2b). The reader displayed in Fig. 2b) contributed 1056 reads and was selected at the median of fit among those in the top decile by read count (*n* = 284 readers), so selection was based here on sample size, rather than on goodness of fit. Across all readers the median threshold is *θ* = 17.41 and the median drift rate *α* = 0.1597, both in units of noise amplitude. Moreover, with 96.8% of readers favouring the variable-start over the constant-start model, the advantage is nearly universal rather than driven by a small subset of readers. Furthermore, since no pooling of individuals occurs, this advantage cannot be attributed to the aggregation of heterogeneous individual responses.

Considering this individual perspective, and assuming each reader’s fitted density describes that reader adequately, then the read-weighted average of those densities must reproduce the pooled distribution:

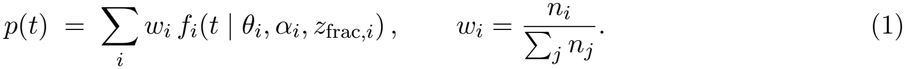

The resulting curve has no free parameter at the population level, so Fig. 2c) is a prediction whereas Fig. 2a) is a fit. Its constant-start counterpart was obtained by refitting every reader under the constraint *z*_frac_ = 0, rather than simply setting the starting point of the variable-start fit to zero, which would not give the constant-start model.This is a stronger test than the pooled comparison in Fig. 2a), where the parameters were free to adapt to the aggregate shape itself. It is not, however, decisive and misfits in individual densities can offset one another in the average, so agreement at the population level is necessary but not sufficient for the individual fits to be correct.

The three views agree that the starting point varies. Whether that confirms evidence accumulation is a separate question, since both candidates are drift-diffusion models, and we addressed it by comparing the same fits against a wider set, six other drift-diffusion models with time-varying drift and seven descriptive families with no generative content, at all three levels of aggregation. We revisit the same three views against this wider set.

For the pooled view, the variable-start model gives the closest match to the empirical distribution of any candidate on both the Kolmogorov–Smirnov and Wasserstein-1 distances tests, at every analysis window (Supplementary Fig. S4). It does not, however, attain the highest pooled held-out likelihood and a seven-parameter drift-diffusion model with time-varying drift exceeds it on that measure at all three windows, without matching it on the Wasserstein-1 distance at any of them, consistent with additional drift parameters absorbing variation that arises between readers rather than within them. For the individual view, the variable-start model is preferred over every other drift-diffusion model (in particular, over the constant-start model for 97% of readers and over the previously mentioned seven-parameter time-varying accumulator for 67%) while three descriptive families are indistinguishable from it, a Gamma, a Weibull, and a power law with exponential cutoff, all within two points of chance (Supplementary Fig. S5).

For the mixture view, which has no free parameter at the population level, the variable-start model ranks first on all three measures. It exceeds every other drift-diffusion model, the seven-parameter time-varying one included (rank-biserial +0.33 on held-out likelihood, Wilcoxon signed-rank *p* < 0.001 after Holm correction), while the same three descriptive families remain indistinguishable from it (Supplementary Fig. S6).

On the other hand, the Gamma distribution, a two-parameter density with no generative content, predicts these reading times as well as the accumulation model does (as do the Weibull or the power law with exponential cutoff). But at the resolution these data provide, goodness of fit cannot separate an accumulation account from a sufficiently flexible description. In fact, what distinguishes them is not how well they describe the observed distribution but the explanatory content that survives the fit. The parameters obtained from the variable-start model are claims about a reader (e.g., how much evidence they require, and how fast an article supplies it), and both are recoverable properties of individual readers rather than artifacts of estimation (Supplementary Fig. S7). These reader-level parameters also have implications for other aspects of the same reader’s behaviour and we test whether the individual differences captured by the variable-start model also explain differences in navigation behaviour in later sections. A fitted Gamma distribution, by contrast, provides no equivalent behavioural predictions (its parameters describe the shape of the reading-time distribution but do not characterise underlying reader-level processes).

Furthermore, one can observe that these conclusions do not depend on the analysis window and repeating the analysis at *t*_max_ = 100, 300 and 900 s leaves the ordering unchanged (Supplementary Figs. S4, S5 and S6). The one exception is the mixture view at *t*_max_ = 100 s, where the descriptive families overtake the variable-start model, but it should also be noted this is the least informative window, since truncation at 100 s places a substantial part of many readers’ fitted densities outside the observed range, and was used as a limiting case.

### 2.3 Evidence accumulation model does not account for the switching time between articles

If the decision to leave an article can be interpreted as an accumulation to a threshold, the natural next question is whether the decision to go and open the next article can be accounted for using a similar process. The two are different problems within the foraging framework (one concerns the allocation of time within a patch, the patch problem, and the other the evaluation of cues before moving to the next patch, the scent problem), but the formalism is not a priori necessarily confined to the first, and whether it transfers to the second is an empirical question

The interval between leaving one article and opening the next is equivalent to the travel time, *τ* , within the foraging framework. The data format allows us to observe the transition rather than infer it. A reader leaving an article either opens another one directly or returns to a navigational page first. We count only returns to the front page as feed visits and exclude transitions passing through a section page, which only shows articles from that single section, so a return there cannot expose the reader to alternatives from elsewhere and would attenuate the topic contrast rather than inform it (these represent 9.1% of all transitions within the data we are using). For returns to the front page, the tracker records the time spent there (Fig. 3a). After this exclusion, 8,703,877 within-session transitions from 682522 readers remain, of which 59.0% pass through the feed and 41.0% are direct. The feed visit represents a search episode in which readers browse for their next article and we treat its duration as the travel time, *τ*.

**Figure 3:**
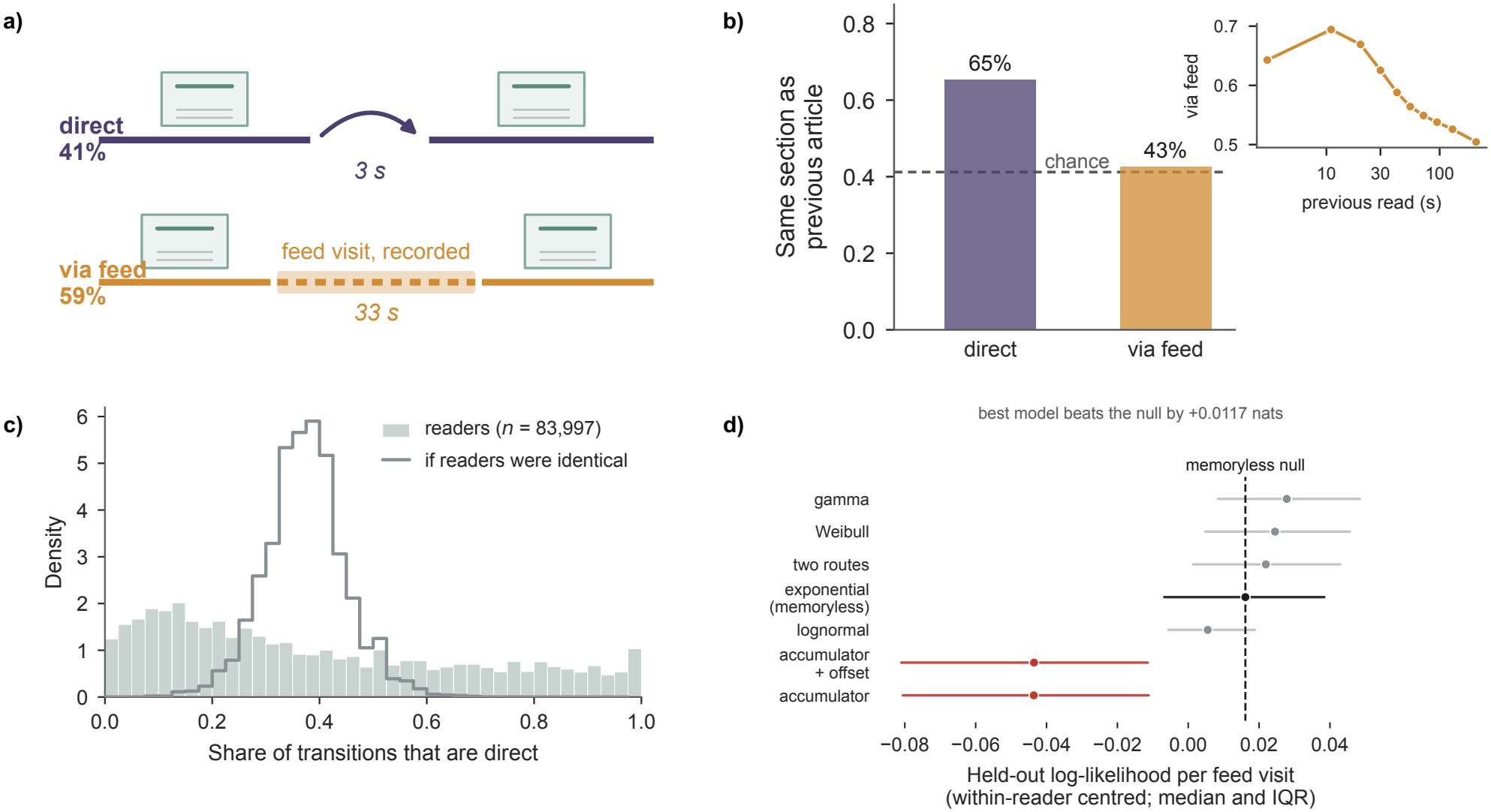
Time to next information patch. Transitions are consecutive article visits within a reading session. **(a)** Schematic of the two routes an individual could take after leaving a patch, a direct transition to another article (purple, 41% of transitions, median gap 3 s), or a transition through the front page (orange, 59%, median gap 33 s), during which the tracker records the time spent on the feed. **(b)** Probability of transitioning to an article with the same section as the previous for both direct (purple) and via feed transitions (orange). The dashed line marks the chance level obtained by permuting each reader’s destination sections within their own reading history, so that their section composition is preserved and only the ordering is destroyed. Inset represents the percentage of transitions passing through the feed as a function of the reading time of the preceding article, binned by decile displaying a decreasing behaviour. The horizontal axis is logarithmic **(c)** Distribution across readers of the share of transitions that are direct, for readers contributing at least 20 transitions (green, n = 83,997), against the distribution expected if readers were identical (grey), obtained by drawing each reader’s count of direct transitions from a binomial with their own number of transitions and the overall direct rate. **(d)** Held-out log-likelihood per feed visit for seven candidate distributions fitted to the feed-dwell times, each estimated on a random half of a reader’s visits and evaluated on the other half. Values are centred within reader and shown as medians with interquartile ranges. The dashed line marks the exponential (memoryless) model and drift-diffusion models are shown in red. Higher values indicate better prediction.

The two routes differ in how often they lead to an article from the same section. Direct transitions lead to an article from the same section as the previous one for 65.4% of 3571698 cases, while feed transitions do so for 42.7% of 5132179, (Fig. 3b). The within-reader permutation baseline is 41.0% (obtained by shuffling each reader’s destinations within their own reading history so that their section composition is preserved and only the ordering destroyed). The direct route therefore exceeds chance by 24.4 percentage points, while the feed route does so by 1.7. The contrast also holds within readers. Among 43089 readers contributing at least ten transitions of each kind, the median difference in same-section rate between the direct and feed routes is 25.1 percentage points, and 90.3% of readers show the difference in the same direction (Wilcoxon signed-rank, *p* < 0.001; matched-pairs rank-biserial *r* = 0.813). The routes also differ in duration, with the direct click possessing a median of 3 seconds against 33 seconds through the feed.

Route, duration and topic therefore covary. One interpretation is that a reader either continues with similar content, in which case the next article is readily accessible and topically adjacent, or seeks different content, which requires returning to the feed and takes longer. The route also responds to what has just been read. Partitioning transitions by the length of the preceding article, the share passing through the feed peaks at 69.4% after reads of about 11 s and declines steadily to 50.4% after the longest decile (median 210 s). Reads shorter than a few seconds are the exception, at 64.3%, consistent with their being misclicks rather than rejections. Generally, a reader who has rejected an article after a short read is more likely to return to the feed than one who has finished a long one, so the choice is conditioned on the information patch they just left and not only on what the reader wants next.

The share of transitions that are direct varies far more across readers than sampling alone would produce (Fig. 3c). Among 83997 readers with at least 20 transitions, the median reader contributes with 44 transitions. To establish what sampling alone would give, we drew for each reader a new count of direct transitions from a binomial with their own number of transitions and the mean per-reader direct rate of 0.373 as probability (the distribution expected if readers were identical). At the median count that null (in grey) has a standard deviation of 0.073, while the observed distribution (in green) is far wider and strongly bimodal, with mass at both extremes. The comparison is a null rather than a model and it shows that the spread is not sampling noise, which licenses treating route preference as a property of readers (but it does not necessarily establish that readers fall into types). We modeled the time spent on the feed before opening the next article as a first-passage problem, in the same manner as reading time, and compared it against six alternatives, chosen due to the format of the distribution, which is concentrated at short durations. Using 20535403 feed visits from 84756 readers contributing at least 100 such visits, we fitted each candidate model to a random half of each reader’s visits and scored on the other.

Scored on held-out visits, the inverse-Gaussian first-passage density obtains −4.3227 nats per visit and its shifted variant −4.3225, against −4.2510 for a Gamma distribution, −4.2579 for a two-component exponential mixture and −4.2675 for a single exponential (Fig. 3d). The two drift-diffusion models are the only candidates whose within-reader-centred intervals lie completely below the memoryless null. Whatever governs the feed dwell time, it does not have the shape of a constant-start first passage to a threshold. Within the models we used to fit, what does describe it is barely distinguishable from a memoryless process. The Gamma distribution exceeds the exponential by a median of 0.0041 nats per visit, for 66% of readers, while the drift-diffusion models lie 0.07 below it. The exponential is the single best candidate for 14% of readers, more often than both drift-diffusion models combined.

An accumulation process describes the decision to leave an information patch, which is the patch problem in the foraging framework, but it does not describe the interval that follows, which belongs to the scent problem. Foraging models themselves treat that interval as a parameter rather than as a decision Charnov [1976], Davidson and El Hady [2019], and our result is consistent with that treatment. A plausible explanation is that feed dwell is dominated by the mechanics of scanning a list rather than by a decision making process with a termination rule (e.g. a reader scrolls until something catches their attention, and the time this takes depends on what the feed contains). One cannot test this hypothesis with the data we are using, since the set of headlines visible at each moment is not recoverable from server logs (the same limitation that prevents us from modeling the diet problem of selecting an article among available alternatives). What the comparison is able to establish is that the interval between patches is not a constant-start first passage to a threshold, and what describes it best improves on a memoryless process by less than one hundredth of a nat.

### 2.4 Patience and assessment floor predict different behaviours

The previous section established that accumulation to a threshold describes the time spent within an information patch, but not the interval between patches. It leaves open however whether the model parameters can tell us some about the behavioral heterogeneity of the readers.

The model assigns to each reader a patience and an assessment floor. Whether those numbers describe the reader or only the fit does it has to be settled before anything is analyzed on them. In order to do this , we simulated realistic readers to reproduce the dynamics of new reading observed in the data. On simulated readers the estimator recovers patience and drift rate with rank correlations of 0.972 and 0.899 (Supplementary Figs. S7**a** and **b**), respectively. On real readers, fitted independently to two disjoint halves of their own reads, the same parameters reproduce at 0.887 and 0.780 (Supplementary Figs. S8**a** and **b**). Patience and the drift rate can therefore be considered stable properties of a reader. The start fraction replicates less closely, and the consequences of that for the assessment floor are laid out in the Supplementary Information.

Across readers the fitted assessment floor (*d*_*min*_) is bimodal, with one mode around *d*_*min*_ = 3 and one close to *d*_*min*_ = 0, and almost nothing between them (Fig. 4a). A Gaussian mixture on the fitted parameters (Methods) recovers that exact division without being given the assessment floor. The number of components was chosen by split-half reproducibility rather than by the Bayesian information criterion. In fact, BIC reaches its lowest value at *k* = 7, while the adjusted Rand index between the labeling of the two half-sample mixtures assigns to the full sample peaks at 0.96 for *k* = 3 (Supplementary Figs. S10**a** and **b**).

**Figure 4:**
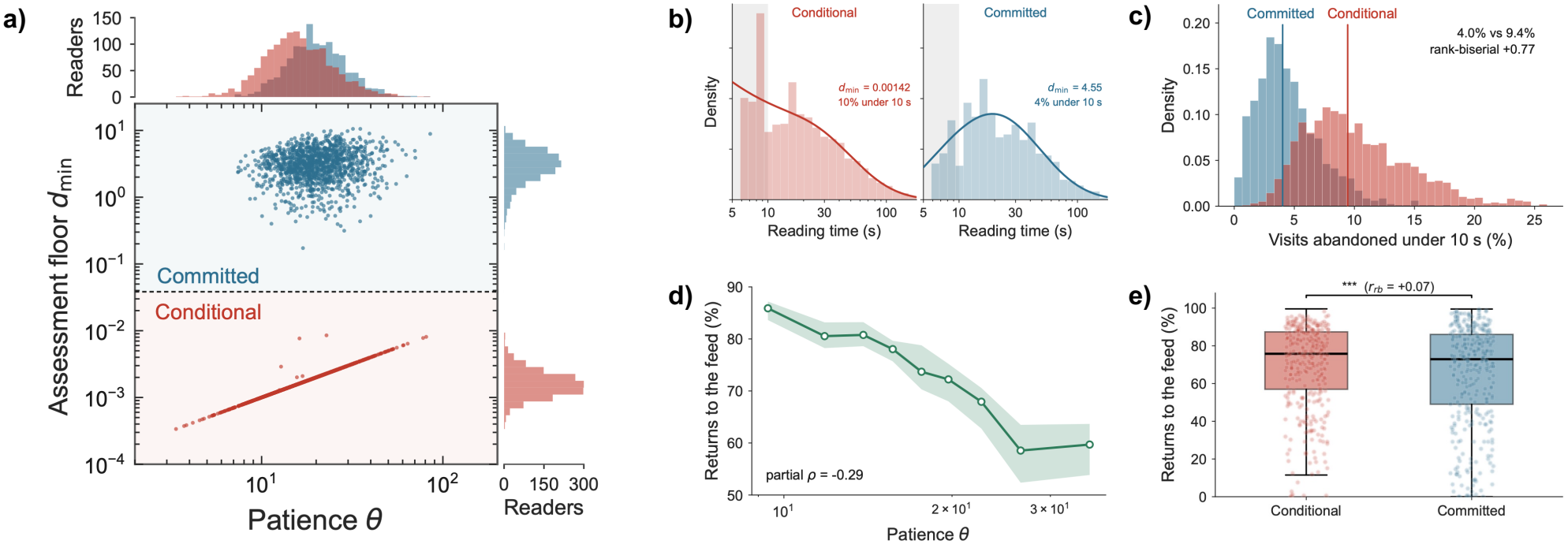
Clustering of users reflect the existence of different behavioural patterns. **(a)** Scatter plot of fitted parameters, assessment floor (*d*_*min*_) and patience (*θ*), of the variable-start model for each reader, with marginal distributions above and to the right. Gaussian mixture on the log-scaled parameters separates 2,706 classified readers into a committed cluster (blue, *n* = 1, 235) and a conditional cluster (red, *n* = 1, 471) and readers the mixture could not place on one side of the division are excluded (Methods). The dashed line marks the boundary between clusters. The fitted assessment floor is bimodal across readers (Hartigan’s dip test on log_10_ *d*_min_, *D* = 0.151, *p* < 10^−3^), while patience is unimodal (Hartigan’s dip test on log_10_ *θ, D* = 0.005, *p* = 0.98), so the clusters are separated by the floor and not by patience. **(b)** Reading times of one reader from each cluster, conditional (red, left) and committed (blue, right), as histograms with their fitted densities. The readers are matched on drift rate and read count (*α* = 0.163 and 0.161, with *n* = 879 and 852 reads). They differ in the assessment floor (*d*_min_ = 0.00142 against 4.55) and in the share of visits ending within 10 s (10% against 4%). **(c)** Distribution across readers of the percentage of visits ending within 10 s, for the committed (blue) and conditional (red) clusters. Vertical lines mark the cluster medians, 9.4% and 4.0% respectively (rank-biserial correlation +0.77). **(d)** Median share of transitions passing through the front page, by quintile of patience. The shaded band is the bootstrap 95% confidence interval of the median. The partial rank correlation between patience and this share, controlling for the assessment floor, is −0.29. **(e)** Share of transitions passing through the front page, for the committed (blue) and conditional (red) clusters. Boxes span the interquartile range with the median marked, whiskers extend to 1.5 times that range, and points are a random sample of readers. The bracket reports a Mann-Whitney test and the rank-biserial correlation (^***^*P* < 0.001).

Two of the three components have median assessment floors of 0.0016 and 0.0027 and are separated by patience rather than by the value of the floor, with the third having a median of 3.12. Readers that the mixture could not place on one side of the division were excluded (Methods), leaving 2,706 classified readers: 54.4% whose fitted assessment floor is close to zero (red), which we term conditional, and 45.6% with a substantial floor (blue), committed (Fig. 4a), right histogram).

The two groups occupy almost the same range of patience, *θ*, with their marginal distributions overlapping across it (Fig. 4a), top histogram). The fitted assessment floor is bimodal across readers (Hartigan’s dip test on log_10_ *d*_min_, *D* = 0.151, *p* < 10^−3^), while patience is unimodal (Hartigan’s dip test on log_10_ *θ, D* = 0.005, *p* = 0.98). The clusters are therefore separated by the assessment floor, not by a difference in patience.

Two readers matched on drift rate and read count (*α* = 0.163 and 0.161; *n* = 879 and 852 reads) (Methods) differ in the shape of their reading-time distributions (Fig. 4b). The conditional reader’s fitted density is highest at the shortest observable times and falls monotonically from there, with 10% of visits ending within 10 s, while the committed reader’s rises from 5 to about 20 s before decreasing in a similar fashion, with 4% ending within 10 s. The assessment floor is what separates them (*d*_min_ = 0.00142 against 4.55) and, since this variable is what sets the minimum distance any accumulation process must travel, it constrains the shortest visits and has a much smaller effect on the rest of the distribution.

This observation is not exclusive to these two readers. Pooled across readers, the two clusters’ readingtime distributions differ almost entirely below 10 s and tend to converge above roughly 30 s (Supplementary Fig. S11f), as a difference in the assessment floor predicts and a difference in patience would not. At the reader level, conditional readers end 9.4% of their visits within 10 s compared with 4.0% of visits by committed readers, yielding a rank-biserial correlation of +0.77 (Fig. 4c). Importantly, this measure is calculated directly from observed reads and was not used to fit the model, so the separation between clusters reflects a genuine difference in reader behaviour, rather than merely a difference in the parameter space.

The second behaviour comes from the navigation analysis of Fig. 3 and is computed from the share of a reader’s transitions between articles that pass through the front page (via feed). Across the range of patience, the median falls from 86% to 59% (Fig. 4d, partial rank correlation of −0.29, controlling for the assessment floor). Readers who require more evidence before leaving an article take a more direct route to the next one. The previous section showed that a first-passage model fails to account for the traveling between information patches, but patience, estimated from behaviour within a patch, nonetheless predicts the route they take to the next one. Divided by the assessment floor rather than by patience, the same behaviour barely separates, with 75.8% of transitions through the feed against 72.9% and a rank-biserial correlation of +0.07 (Fig. 4e). Patience and the assessment floor correlate at 0.564 across readers, so a marginal comparison cannot attribute either behaviour to one of them alone (Supplementary Table S5). Partial rank correlations, each controlling for the other parameter, separate them (Table 1).

**Table 1:** Partial rank correlations between the fitted parameters and two behaviours (navigation via feed and visits under 10 s). Each controls for the other parameter only, since the behaviours are consequences of the parameters rather than competing predictors, so conditioning on one while measuring the other would remove part of the path under test.

|  | navigation<br>(via feed) | abandonment<br>(visits under 10 s) |
|---|---|---|
| patience *θ* | −0.291 | −0.412 |
| assessment floor *d*<sub>min</sub> | +0.017 | −0.716 |

The assessment floor strongly predicts information-patch abandonment but provides no evidence of a relationship with navigation (its partial correlation with transitions via the feed is only +0.017, which is not only negligible but also opposite in sign to the expected relationship). Patience predicts both, though more weakly than the assessment floor on abandonment (−0.412 against −0.716). This asymmetry is precisely what distinguishes a parameter that corresponds to a property of the reader from one that merely improves the fit. A quantity correlated with every behaviour measured identifies nothing in particular, but one that predicts the behaviour it should and fails to predict the behaviour it should not is measuring something specific.The assessment floor is defined as the minimum evidence any decision must accumulate. Its effects are concentrated in the shortest visits, which is why it predicts abandonment and not what happens once the article is left.On the other hand, patience is defined as the total evidence required, so it governs how long a visit lasts (and, as the previous section showed, how long the previous read lasted is itself the strongest predictor of whether a reader returns to the feed).

These results position our findings in an information foraging context. Patch exploitation and the travel between patches are ordinarily treated as separate problems, each with its own decision rule Charnov [1976], Davidson and El Hady [2019]. Here the same parameter, *θ*, predicts both. A reader who requires more evidence before abandoning an article also takes a more direct route to the next, and the relation survives while holding the assessment floor constant. The assessment floor, by contrast, predicts only what happens within an article. We take this as evidence that patience is a disposition a reader brings to the session rather than a quantity defined only within an article, though these data cannot establish that directly.

## 3 Discussion

In this study, we applied the information foraging framework to news reading, treating an article as an information patch and the decision to leave it as the outcome of an evidence-accumulation process (a class of models conventionally used in neuroscience to study decision making). The framework distinguishes three problems Pirolli and Card [1999], the patch problem (time allocation within articles), the scent problem (headline evaluation on the front page) and the diet problem (which article is selected). We model the first directly, observe but do not decompose the second, and do not address the third, because the set of alternatives visible at any moment is not recoverable from server logs of the news data we are using in this study.

We have shown that reading times within an article are described by accumulation from a variable starting point, in which a reader arrives already partly informed. In systems and cognitive neuroscience, decision-making has been extensively studied using the concept of evidence accumulation Stone [1960], Laming [1968], Link [1975], Ratcliff and McKoon [2008], Falmagne [1965], Bogacz et al. [2006], Piet et al. [2018]. Evidence accumulation has been implicated for example in social decisions [Krajbich et al., 2012], sensory decisions [Brunton et al., 2013, Hanks et al., 2015], economic decisions [Gluth et al., 2012], memory decisions [Ratcliff, 1978], visual search decisions [Purcell et al., 2010], and value decisions Milosavljevic et al. [2010]. What distinguishes the present application is not the model but the setting and the resolution. In our study, the decisions are unprompted, the alternatives were not selected by an experimenter and the record is long enough per person to estimate a leaving rule for each.

On the other hand, the evidence accumulation process was not able to predict the interval between articles. A constant-start drift-diffusion model is a poor description of the time a reader spends on the feed before opening the next article, and the best of the candidates we compared offers little improvement in holdout likelihood over an exponential distribution, which represents a memoryless process. In foraging terms, the patch problem and the scent problem are different problems, and our results suggest they are not described by the same formalism. Foraging models themselves treat the interval between patches as a parameter rather than as a decision Charnov [1976], Davidson and El Hady [2019], and our result is consistent with that treatment. One plausible explanation is that feed dwell time is governed by the mechanics of scanning a list rather than by a decision process with a termination rule, so that the time taken depends on what the feed contains rather than on what the reader requires. We cannot test this hypothesis with the present data, for the same reason we cannot address the diet problem.

Fitting each reader separately revealed that readers are not distributed smoothly in the parameter space. A Gaussian mixture model separates readers into two groups that differ in their assessment floors, even though this quantity was not used in the clustering process. For one group, the upper bound of the initial evidence distribution falls below the decision threshold, so the model requires a minimum amount of evidence that must be accumulated before a decision can be reached. For another group, it requires none the data can detect. Readers in this second group abandon articles within 10 seconds of arrival at more than twice the rate observed in the first group. Thus, the groups are distinguished not only by their fitted model parameters but also by differences in independently observed reading behaviour. This suggests that the clustering captures meaningful differences between readers, rather than simply partitioning the parameter space.

The assessment floor and patience capture distinct aspects of reader behaviour. Holding the assessment floor constant, patience predicts both how often a reader abandons an article and whether they return to the front page before opening the next. Holding patience constant, the assessment floor predicts abandonment strongly and does not predict navigation between articles. Note that a parameter that correlates with every measured behaviour tells us little about what it specifically captures, while one that predicts the behaviours it is expected to predict, but not those it is not, provides evidence that it captures a specific aspect of reader behaviour. This asymmetry also argues against the alternative that both parameters reflect the same underlying reader disposition. If they did, we would expect both parameters to predict both behaviours. Instead, their distinct predictive patterns suggest that they capture two different properties of readers rather than measuring the same property twice.

In the information foraging framework, patch exploitation and the traveling between patches can be conceptualized as separate problems, each with its own decision rule. Here one parameter reaches both. A reader who requires more evidence before abandoning an article also takes a more direct route to the next, and the relation survives holding the assessment floor constant. The assessment floor, by contrast, predicts only what happens within an article, which is what a quantity, setting the minimum evidence any decision requires, should do. We take this as evidence that patience is a disposition a reader brings to the session rather than a quantity defined only within an article.

Our study have some limitations. First, the data are from a single regional newspaper, in one language, over ten weeks, so we cannot say whether the behavioral categories we report reflects something general about readers or something about this outlet and its particular audience. Second, the readers analysed are by construction the most active on the site, since fitting three parameters per reader requires a long record, and they are therefore not representative of the totality of its visitors. Third, the assessment floor is identified in ordering rather than in value for readers near the parameter bound, so the conditional group is defined by the absence of a detectable floor rather than by an estimate of a small one. A second dataset with per-article engagement times and comparable per-reader depth would test whether the division and the dissociation replicate.

Several concrete next steps could build up on this study. A dataset with richer metadata would allow the scent and diet problems to be addressed rather than observed, giving a deeper understanding of digital news consumption. This approach can also be extended to platforms such as Reddit, X, Bluesky and TikTok Pirolli [2020], Drias and Pasi [2020], where the difficulty would be establishing the nature and amount of information a user is sampling and evaluating when interacting with a post on any of these platforms Roozenbeek et al. [2025]. The model could also be extended to include social information Pirolli [2009], Luca et al. [2009], Pirolli [2008], since time spent on an article may be shaped by knowing that others have read or shared it, a question better addressed by experiments in which readers are asked about such influences Metzler and Garcia [2024]. Comparing the models we developed in this study to a reinforcement learning algorithm might give us insights into potential learning processes involved in news consumption, which could equally be formalised as Bayesian inference Kilpatrick and El Hady [2026], with reading shaping a reader’s posterior probabilities making them more likely to read some articles rather than others and so differentiating readers over time. The latent variables of a drift-diffusion model may also be used to study how misinformation spreads, for example by asking which of them communicate confidence in information.

More broadly, our study further highlights the idea that foraging is a general adaptive process operating over internal as well as external resources Hills et al. [2008, 2012], Hills and Butterfill [2015], Todd and Hills [2020], Safavi and Dayan [2022], and contributes to the increased interest in casting information as a key currency in defining the ecological, neural and behavioural constraints animals and humans face in their environments, an area of inquiry termed information ecology, which studies how individuals use information to enhance fitness Metzler and Garcia [2024].

## 4 Materials and methods

### 4.1 Data description

We analyzed user activity information from *Adresseavisen*, a regional newspaper published in Trondheim, Norway, released as the Adressa dataset which was originally prepared as part of the RecTech project on recommendation technology Gulla et al. [2017]. These activity logs provided fine-grained information about users’ interactions and navigation behavior, enabling the reconstruction of digital movement alongside identification of information foraging events.

The data were collected for 10 weeks (from 1 January to 31 March 2017) and are made up of 5.83 × 10^7^ page-visit records from approximately 3.1 × 10^6^ readers. Each record contains a pseudonymous reader identifier, the visited URL, a Unix timestamp of the page request, boolean flags marking session boundaries, and `activeTime` (the seconds of active engagement on the page). Nevertheless, not every record corresponds to an article read. We therefore classified pages according to their URL patterns. Article pages (which correspond to the information patches in the foraging framework) were identified by the terminal pattern (for example, `.ece` or `.html`). The front page (`adressa.no`) is where headlines are assessed. Section pages (for example, `/nyheter/` or `/100sport/`) are intermediate navigation surfaces. Across the dataset, 34.3% of records are articles, 56.6% are front-page visits, and 9.0% are sections or other pages. In the information foraging framework of Pirolli and Card Pirolli and Card [1999], this classification maps onto their three model types. In this paper, we model the patch problem (time allocation within articles) directly, we observe but do not decompose the scent problem (headline evaluation on the front page) and we do not model the diet problem (which article is selected), because the set of alternatives available at each moment is not recoverable from server logs.

### 4.2 Reconstruction of digital trajectories

To recover the reading sequences, we reconstructed per-reader trajectories from the raw event log by segmenting it into sessions, restricting it to article pages, and merging fragmented visits. In fact, the tracker reports events, not reading episodes per se, and these three reconstruction steps separate the two. In practice, for each reader, events were ordered by timestamp and a new session was opened whenever the interval between the end of one event and the start of the next exceeded 300 s, or whenever the tracker’s own session flag was set. This 300 s threshold is a convention rather than a measurement and formalizes the assumption that a reading session remains engaging until a pause of five minutes or longer occurs, at which point a new session begins (we therefore verified that the results are insensitive to this threshold; see Fig. S4, S5 and S6).For every event, we also computed the time elapsed since the previous article was exited, the travel time (*τ*).

Events were restricted to article pages (the patches of the information foraging framework) identified by URLs ending in `.ece` or `.html`. Front pages, section pages and other navigational pages were therefore excluded. Moreover, consecutive events sharing a URL and a session, and separated by 30 s or less, were merged into a single visit whose engagement time is the sum of the fragments and whose remaining fields are taken from the first. This step merged 329382 records and leaves 19688444 patch visits. Readers contributing more than 500 patch visits were retained, yielding 2820 readers and approximately 3.5 million visits. This threshold reflects an estimation requirement. Because each reader is fitted separately, a sufficiently large number of reads is needed to reliably estimate the model’s three parameters. Note however that the retained readers are, by construction, the most active on the site, so the sample is not representative of all readers, and no inference about the wider population can be made from it.

### 4.3 Analytical model

Within each patch visit, a decision variable *X*(*t*) accumulates noisy evidence about the article’s value and the reader then leaves at the first time *T* for which *X*(*T*) = *θ*. The evidence evolves according to:

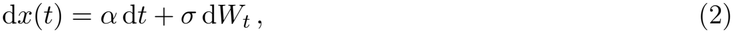

Because only the leaving action is modeled, the process has a single absorbing boundary rather than the two-boundary form familiar from choice tasks Ratcliff and McKoon [2008]. This is a deliberate reduction, considering that a second boundary would represent a competing terminal act, and none is observed in the dataset we are using. On each visit the starting point is drawn afresh, *X*(0) = *z* ∼ *U* (0, *z*_frac_*θ*), so that the remaining evidence to be accumulated, *d* = *θ*− *z*, is uniform on [*d*_min_, *θ*] with *d*_min_ = *θ*(1− *z*_frac_). Across-trial variability in the starting point is a standard feature of diffusion models Laming [1968], Ratcliff and Rouder [1998], Boehm et al. [2018], where it is conventionally interpreted as response bias, a prior tendency toward one response over another. Here it represents evidence already sampled before the accumulation window opens, which is the natural diffusion-model expression of pre-sampled evidence.

For a fixed starting point, the first-passage density of Eq. (2) to a boundary at distance *d* is the following inverse Gaussian:

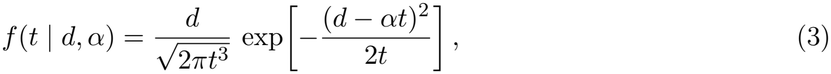

where *d* = *θ* in the constant-start variant. Under a variable-start *d* is uniform on [*d*_min_, *θ*], and averaging Eq. (3) over that distribution admits the closed form below, which is evaluated directly rather than by quadrature. By writing 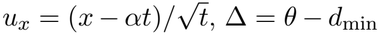 and by denoting the standard normal distribution and density as Φ and *ϕ*, we obtain

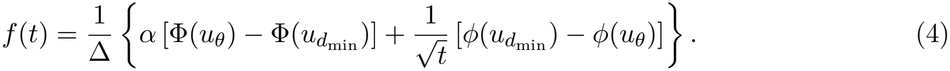

The diffusion coefficient is fixed at *σ* ≡ 1 (the first-passage density depends on *θ* and *σ* only through their ratio, so one must be constrained). Consequently, *θ* and *α* are expressed in units of noise amplitudes rather than in seconds.

Reads were restricted to the window [*T*_min_, *t*_max_). The lower bound was fixed at *T*_min_ = 5 s to remove accidental clicks and page bounces. The upper bound was varied over *t*_max_ ∈ {100, 300, 900}s, with 300 s reported in the main text. The other time intervals are used to establish that the conclusions do not depend on the particular choice of the value of *t*_max_ (see Fig. S4, S5 and S6). Because reads are observed only inside this window, the density is conditioned on it, and a uniform contaminant of weight *π* = 0.02 is added so that a single mistracked read cannot dominate the sum:

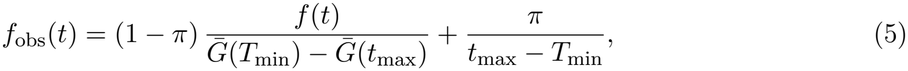

where 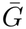 is the survival function of the corresponding variant 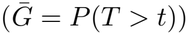. Scoring a model over a window different from that used for fitting can bias model comparisons, with the magnitude of the bias depending on the probability mass falling outside the evaluation window. The contaminant weight was fixed rather than estimated.

Parameters were estimated for each reader separately by maximum likelihood using L-BFGS-B under box constraints:

- *θ* ∈ [0.01, 500],
- *α* ∈ [0.001, 10],
- *z*_frac_ ∈ [0.01, 0.9999].

The upper bound on *z*_frac_ is set below 1 because by definition one has *d*_max_ > *d*_min_. *θ* and *z*_frac_ trade off along a direction of nearly constant effective distance and a single start is not reliable on such a surface, so optimisation was repeated from six starting points, spread across the range of *z*_frac_. Note that readers were estimated separately rather than hierarchically because shrinkage toward a group mean would attenuate precisely the between-reader variation the analysis is designed to characterise.

### 4.4 Time between information patches

Each logged page view was assigned to one of three kinds. Articles were identified by URLs ending in `.ece` or `.html`, (the patterns the site uses for editorial content, 34.3% of records). The front page was the bare domain, https://www.adressa.no (56.6%). All remaining on-site pages were classified as navigational (e.g. section fronts, sub-section listings, tag pages and search results).

A transition is a pair of consecutive article views by the same reader within a session. The interval between them is the travel time, *τ* , computed as the timestamp of the second article minus the sum of the first article’s timestamp and its engagement time. Pairs were retained if they fell within the same session, had 0 ≤*τ* ≤600 s, and had no navigational page between them. The last rule follows from what the two routes are meant to contrast. A reader returning to the front page is exposed to articles from every section and may switch topic (while section listings shows articles from one section only, so a return there cannot produce the comparison the analysis makes).Simultaneously, considering such a transition as direct one between articles would attribute to an in-article link a choice the reader made from a list.

Transitions passing through a navigational page are therefore excluded rather than assigned to either route. A transition was therefore classified as *via feed* if at least one front-page passage occurred between the two articles, and *direct* if nothing fell between them. A transition was considered samesection if the two articles shared a section. The observed percentage of this quantity is not interpretable on its own, because a reader who reads predominantly within one section will stay within it by construction, so the baseline was obtained by permutation. Each reader’s sequence of destination sections was shuffled within their own reading history, preserving their section composition and destroying only the ordering. Averaging the recomputed rate over *n* = 25 permutations gives a chance level of 41.0%.

For each route type, direct transitions and feed transitions, we calculated the same-section rate. Similarly, for each reader, we then calculated the same-section rate separately for each transition type and computed the within-reader difference between the two rates. We tested these paired differences across the 43089 readers with at least ten transitions of each type using a Wilcoxon signed-rank test, with the matched-pairs rank-biserial correlation as the effect size.

Furthermore, for each reader with at least 20 transitions, we recorded the total number of transitions and the number that were direct (using counts rather than rates because the sampling variability of a rate depends on the underlying count). To obtain the expected distribution under reader homogeneity, we simulated the number of direct transitions for each reader using a binomial distribution. The number of trials was set to that reader’s total number of transitions, and the success probability was set to the overall direct transition rate across all readers. We then compared this expected distribution with the empirical distribution of direct-transition rates.

The feed dwell is the engagement time recorded on a front-page interaction, restricted to the [1, 600] s interval and with longer visits being treated as interruptions rather than search. Only readers contributing at least 100 of such visits were retained.

Seven candidate distributions were compared:

- Exponential distribution (the memoryless null)
- Log-normal distribution
- Weibull distribution
- Gamma distribution
- Two-component exponential mixture
- Constant-start drift-diffusion model
- Constant-start drift-diffusion model with offset.

The candidate set spans both mechanistic and descriptive accounts, so that a descriptive family winning would indicate that the data do not support a decision process rather than that the wrong mechanism was tried. Each candidate was fitted by a maximum likelihood routine from three starting points under box constraints, on a random half of each reader’s visits, and scored on the other half.

### 4.5 Model Parameters, behavioural patterns and individual differences

Each reader contributes a set of fitted parameters from the variable-start model and two behavioural quantities obtained from the data. The first is the share of a reader’s visits ending within 10 s, counted over the reads inside the [5, 300) s analysis window so that the quantity refers to the same observations the model was fitted to. The contrast between clusters is unchanged if the count is taken over all of a reader’s reads instead (16.5% against 8.0%, compared with 9.4% against 4.0% within the window). The second is browsing propensity, the share of a reader’s article-to-article transitions via feed (i.e. transitions that pass through the front page), taken from the navigation analysis described above. ‘ Readers contributing fewer than 20 transitions are left out of the browsing analyses rather than assigned a rate (at that count the sampling noise on a rate is comparable to the between-reader spread the measure is intended to resolve).

Readers were clustered on the fitted parameter (*θ, α*_0_ and *z*_frac_). *θ* and *α*_0_ were log-transformed before standardisation, so that equal ratios correspond to equal distances. *z*_frac_ is a ratio bounded on [0, 1] and was left untransformed (a log would compress the values near one, which is where the two groups separate). Remember that, by definition, *d*_min_ = *θ*(1 − *z*_frac_).

Gaussian mixtures with full covariance matrices were fitted for *k* = 1 to 8. The number of components was selected by split-half reproducibility rather than by the Bayesian information criterion. BIC falls steeply over the first few values of *k* and then varies without a clear trend, reaching its lowest value at *k* = 7 (Supplementary Fig. S10). BIC measures how closely the mixture describes the parameter distributions rather than how reproducibly it assigns readers, and, since our aim is to identify groups of readers, the stability of cluster assignments is more important than goodness of fit alone Reproducibility was measured by fitting a mixture to each half of a random split and computing the adjusted Rand index between the two labelings, averaged over ten splits. Both half-sample models label the full sample, so the index compares two labelings of the same readers rather than of disjoint sets, which gives higher values than the usual construction. This procedure selects *k* = 3.A reader whose maximum posterior falls below 0.75 was left unassigned rather than forced to the nearer component (this applies to 0.7% of readers).The partition reported is between the two largest components, with the 94 readers of the third (3% of those with fitted parameters) being excluded alongside the unassigned.

Hartigan’s dip test was applied to log_10_ *d*_min_ and log_10_ *θ* across classified readers, and repeated on all readers with fitted parameters to confirm that any departure from unimodality is a property of the sample rather than of the partition.

The two example readers shown were selected by minimising a cost combining the following requirements:

- each is located near its own cluster’s median short-visit rate, so the pair is representative of the clusters;
- each lies near the 85th percentile of read count within its cluster;
- the second reader is matched to the first on drift rate and read count.

Matching is necessary because an unmatched pair could differ in drift rate or in sample size, and the panel could then display that difference rather than the one in the assessment floor.

Given that patience and assessment floor are correlated across readers, a marginal correlation cannot attribute a behaviour to either alone. Each reported partial is computed from the inverse of the rank correlation matrix of three variables (the two parameters and one behaviour), which is the Spearman analogue of a partial correlation and assumes nothing about functional form. We deliberately do not control for the second behaviour when analysing the first. Both behaviours are considered outcomes of the reader parameters, rather than predictors of each other. Controlling for one behaviour could therefore remove part of the relationship between the reader parameters and the behaviour we want to examine.

Differences between the two clusters in each behaviour (Fig. 4c & e) are reported as rank-biserial correlations derived from the Mann–Whitney *U* statistic, *r*_rb_ = 2*U/*(*n*_1_*n*_2_) − 1. Both behaviours are proportions bounded between 0 and 1 with skewed distributions, so an effect size based on means and standard deviations would be difficult to interpret.Separately, for Fig. 4d), readers were also divided into quintiles of patience and the median browsing rate computed within each quintile, intervals on the quantile medians are bootstrap percentile intervals from 2,000 resamples of the readers in that quantile.

## Acknowledgments

A.E.H. and H.M.B thanks Konstanz School of Collective Behavior where this project began. A.E.H. acknowledges support from the Deutsche Forschungsgemeinschaft (DFG, German Research Foundation) under Germany’s Excellence Strategy, EXC 2117-422037984 and the Human Frontier Science Program (RGP006/2025). Authors H.M.B. and A.E.H. developed the study, derived the results, and wrote the manuscript.

## Supplementary Materials

### Digital news consumption as an information foraging problem

Henrique M. Borges and Ahmed El Hady

In the following supplementary materials, we delineate detailed data analysis, model derivations, statistical analysis and robustness tests complementing the main manuscript. It is important to note that references are listed in the main manuscript; citation numbers continue that list.

## 1 Model

For a fixed starting point, the first-passage density of Eq. (2) to a boundary at distance *d* is the inverse Gaussian of Eq. (3). Under a variable start the remaining distance *d* = *θ* − *z* is uniform on [*d*_min_, *θ*], since *z ∼ U* (0, *z*_frac_*θ*). Given that a linear transformation of a uniform variable keeps it uniform, the reading-time density is the average of Eq. (3) over that interval:

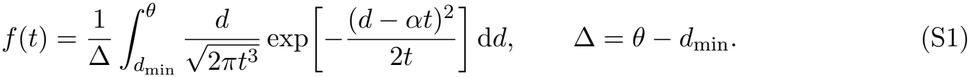

Substituting 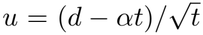, so that 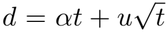 and 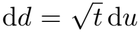, the powers of *t* cancel against the factor *d* in the numerator and the integrand separates into two distinct terms:

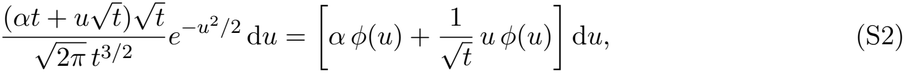

where *ϕ* is the standard normal density.

The first term integrates to the standard normal cumulative distribution function (Φ) and the second term is an exact derivative, since *ϕ*^′^(*u*) = −*uϕ*(*u*), and integrates to *ϕ*(*a*) − *ϕ*(*b*) over [*a, b*]. Evaluating between 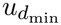 and *u*_*θ*_ gives Eq. (4).

## 2 Estimation

During the estimation process, three methodological choices were made (namely, the analysis window, the bounds placed on the parameters and the weight of the uniform contaminant). Each one of these could, in theory, be responsible for a result attributed to the data. This section reports the practical impact that each choice had in practice.

In theory, a fitted density assigns probability to reading times of any length, including those longer than the window. In practice, a large part of a reader’s implied density falls below *t*_max_, so truncating there is not costly for most readers, but this holds to different degrees for different readers. In this context, Fig.S1 shows precisely how much of each reader’s fitted density falls inside the window. The distribution of retained mass at each window is shown in Fig. S1a). At *t*_max_ = 100 s, the distribution is strongly left-skewed, with a long tail of readers retaining less than half their implied density. As the window widens the distribution concentrates near unity and the 300 s and 900 s distributions overlap substantially, so most of the gain comes from widening the window beyond 100 s rather than from extending it to 900 s. By taking 0.90 as a reporting threshold, one can observe on Fig.S1 b) that the number of readers below this threshold falls from 2647 of 2808 at *t*_max_ = 100 s (94%) to 1576 of 2820 at 300 s (56%) and 1285 at 900 s (45%), with the median retained mass rising from 0.696 to 0.887 and 0.912 across the same windows.

**Figure S1:**
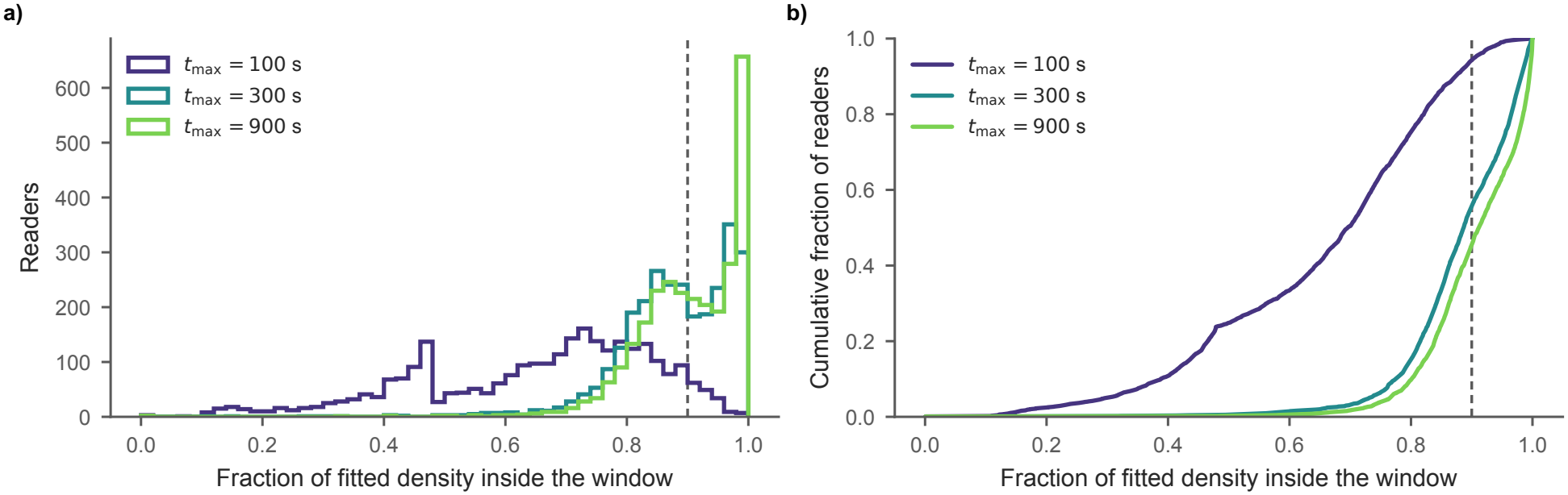
Fraction of each reader’s fitted density falling inside the analysis window. Proportion of each reader’s fitted variable-start density that lies within [5, *t*_max_], computed from their own parameters. **a)** Distribution across readers at each analysis window: *t*_*m*_*ax* = 100*s* (dark blue, n = 2,808 readers), 300*s* (teal, n = 2,832) and 900*s* (green, n = 2,832). **b)** The same distributions as empirical cumulative distribution functions, so that the fraction of readers below any threshold can be read directly. The dashed vertical lines marks a retained mass of 0.90. The numbers of readers below this threshold are 2647 of 2808 readers at 100 s, and 1576 and 1285 of 2820 at 300 and 900 s respectively, with median retained masses of 0.696, 0.887 and 0.912.

In conclusion, at *t*_max_ = 100 s, almost no reader is well characterised, with the median reader retaining barely two thirds of their implied density, and we treat this window as a limiting case rather than as an analysis case. At 300 and 900 s the median reader retains close to nine tenths of their implied density. Nonetheless, 45% of readers remain below the threshold even at 900 s, so widening the window does not remove the censoring but only reduces it. We therefore treat 900 s as another limiting case, rather than as one at which the censoring disappears, and report 300 s in the main text, where the distribution of retained mass is already close to its asymptote. More than half of the fitted start fractions reach the upper bound on *z*_frac_, which potentially invites the objection that the bound was chosen so as to produce this outcome. Fit quality cannot answer it, since a wider bound can never fit worse than a narrower one.

Instead we report three quantities at each combination of bounds. In the first, each reader is fitted twice, on disjoint halves of their own reads, and the two estimates of *θ* are correlated across readers, to check whether the estimates replicate, Fig. S2 a). The second is the held-out log-likelihood, included here for completeness, Fig. S2 b). The third is the proportion of readers reaching the *z*_frac_ bound, Fig. S2 c).

In practice, we refitted 200 readers on both halves of their reads at each of 27 combinations of the three bounds, varying the upper bound on *θ* over {100, 500, 2000}, the lower bound on *α* over {10^−4^, 10^−3^, 10^−2^} and the upper bound on *z*_frac_ over {0.95, 0.99, 0.9999} (Fig. S2). Changing the upper bound on *θ* from 100 to 2000, or the lower bound on *α* from 10^−4^ to 10^−2^, leaves all three quantities practically unchanged, which happens because no reader approaches either bound. However, lowering the upper bound on *z*_frac_ from 0.9999 to 0.95 increases the proportion of readers at the bound from 59% to 62%, as the lower ceiling is more easily obtained, which suggests that the result is not an artifact of a non-restrictive upper bound, considering that it admits fewer readers to the bound than a tight one would. The effect on the other two measures is small in either direction, where across the full range of the upper bound on *z*_frac_ the correlation varies by less than 0.002 and the held-out log-likelihood by less than 0.0005, with the tighter bound giving marginally the higher correlation and the lower likelihood (with neither difference being large enough to favour one value over another).

**Figure S2:**
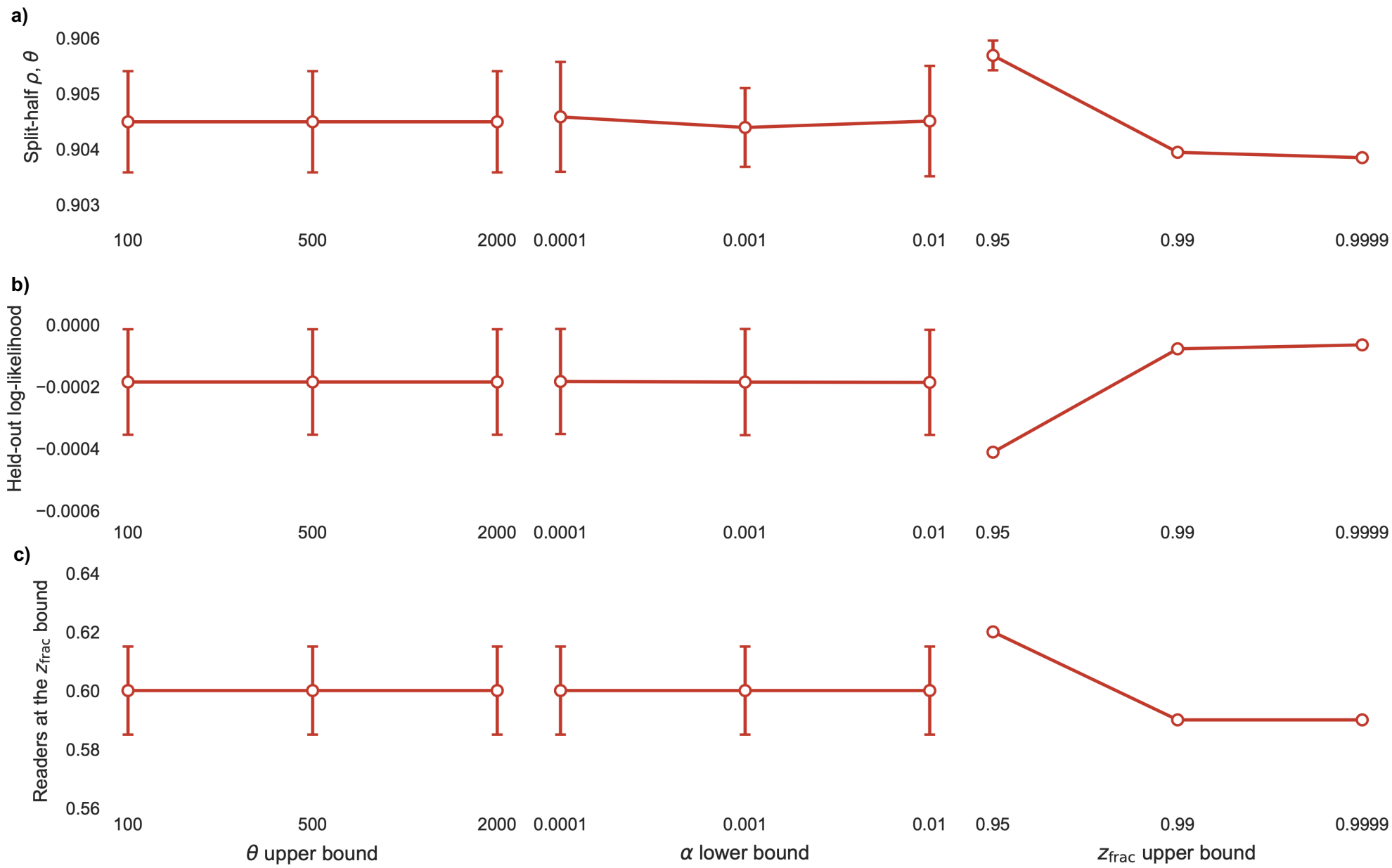
Sensitivity of the estimates to the parameter bounds. 200 readers were refitted on both disjoint halves of their reads for each of the combinations of the three box constraints, the upper bound on *θ* varied over {100, 500, 2000}, the lower bound on *α* over {10^−4^, 10^−3^, 10^−2^}, and the upper bound on *z*_frac_ over {0.95, 0.99, 0.9999}. The analysis window was held fixed at [5, 300) s. Columns correspond to the three bounds, with each point representing the average over the nine combinations of the other two bounds and the error bars indicating the standard deviation across these combinations. All rows share the same vertical axis, such that a flat profile indicates that the bound on the corresponding parameter has no effect on the plotted quantity. **(a)** Spearman correlation between the two half-sample estimates of *θ* across readers. **(b)** Held-out log-likelihood per read, averaged over the two halves. **(c)** Proportion of readers whose fitted *z*_frac_ reaches its upper bound in at least one half (because each reader is fitted twice, this proportion exceeds that obtained from a single fit to the full dataset).

**Figure S3:**
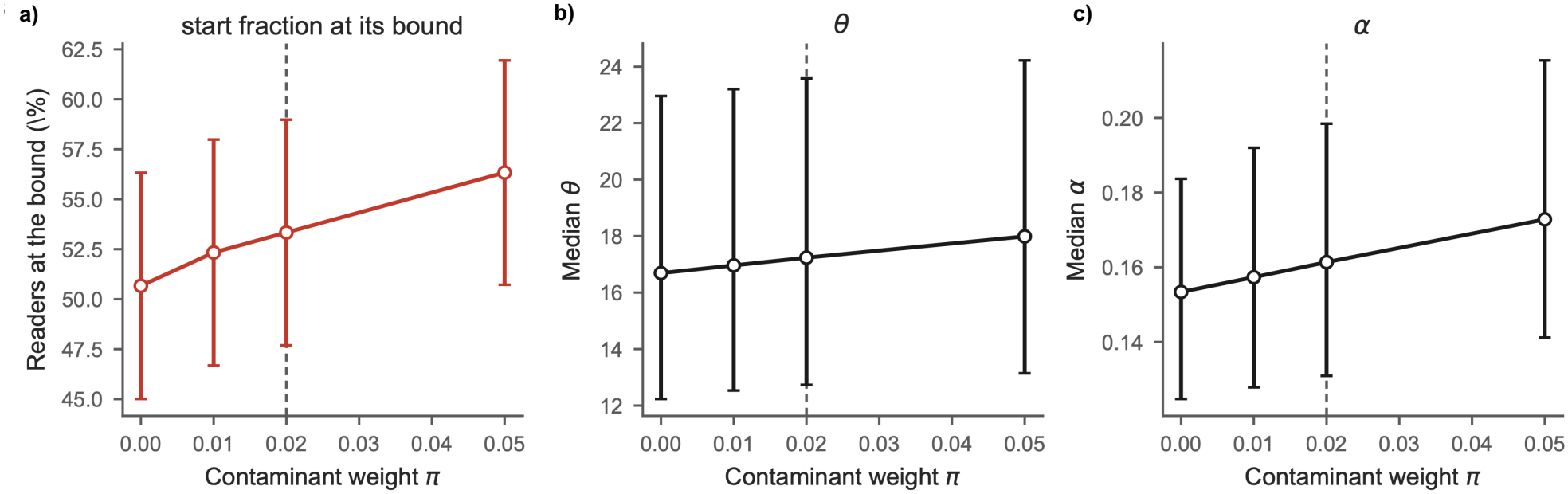
Sensitivity of the estimates to the contaminant weight. 300 readers were refitted at each of four values of the uniform contaminant weight, *π* ∈ {0, 0.01, 0.02, 0.05}, with all other settings held fixed. The analysis window was held fixed at [5, 300) s. The case *π* = 0 corresponds to the no contaminant case. Dashed vertical lines indicate *π* = 0.02, the value used throughout the main analysis. **(a)** Percentage of readers whose fitted start fraction *z*_frac_ reaches its upper bound, with 95% binomial confidence intervals. **(b, c)** Median fitted threshold *θ* and drift rate *α* across readers, respectively, with interquartile ranges. Both parameters are expressed in units of noise amplitude, with the diffusion coefficient fixed at unity.

A uniform contaminant of weight *π* = 0.02 was included so that a single mistracked read could not dominate a reader’s likelihood, and was fixed rather than estimated. Estimated freely, *π* could absorb misfit, and a reader the model is not able to properly describe would be reported as largely noise rather than as a failure of the model.

Refitting 300 readers at *π* ∈ {0, 0.01, 0.02, 0.05} (Fig. S3), the proportion at the bound increases monotonically from 50.7% at *π* = 0 to 52.3%, 53.3% and 56.3%, while the median *θ* rises from 16.69 to 17.99 and the median *α* from 0.1534 to 0.1728 (all changes under 13%).The most informative value is perhaps the endpoint. In fact, with no contaminant at all, 50.7% of readers still reach the bound. The pinning is therefore not produced by the contaminant, and the value adopted in the main analysis sits near the lower end of the range the sweep covers. The proportion is nonetheless not invariant to this choice (it spans 5.6 percentage points across *π* ∈ [0, 0.05], and 1 point between 0.01 and 0.02).

In conclusion, the three analyses demonstrate that the central estimates are insensitive to the procedural choices examined (the analysis window, the parameter bounds and the contaminant weight).

## 3 Model comparison

The variable-start model was compared against thirteen alternatives, chosen to span two kinds of account. Six are *drift-diffusion models*, describing a process that ends when accumulated evidence reaches a threshold, and their parameters refer to quantities a reader could, in principle, possess. Seven are *descriptive families*, so parametric densities with no generative content, which can reproduce a right-skewed distribution without committing to any mechanism that produces it. The drift-diffusion models share the first-passage structure of Eq. (2) and differ in the functional form of the drift. In the *constant-start* model the drift is constant and accumulation begins at zero. The *variable-start* model adds a starting point drawn afresh on each visit. The remaining five allow the drift to vary within a visit:

- *Linear* : *α*(*t*) = *α*_0_ + *βt*.
- *Oscillatory*: 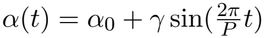.
- *Linear* + *oscillatory*: 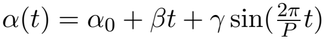.
- *Power* + *oscillatory*: 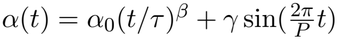.
- *Competing* + *oscillatory*: 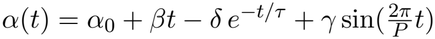.

With constant drift, the expected evidence accumulated by time *t* is *αt*, and Eq. (3) gives the first-passage density exactly. However, when the drift varies within a visit the expected evidence becomes 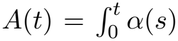 d*s*, and we obtain a density for each time-varying model by substituting *A*(*t*) for *αt* in Eq. (3). The substitution is not exact for the time-varying models, so each density was normalised numerically before scoring. The validity of this approach for inference has recently been questioned **?**.

The descriptive families capture a broad range of temporal patterns through flexible continuous probability distributions. Unlike the drift-diffusion models, these families describe the distribution of observed times directly, each imposing a different functional form:

- *Gamma*: *f* (*t*) ∝ *t*^*k*−1^*e*^−*t/σ*^.
- *Log-normal* : *f* (*t*) ∝ *t*^−1^ exp[−(ln *t* − *µ*)^2^*/*2*s*^2^].
- *Weibull* : *f* (*t*) ∝ *t*^*k*−1^ exp[−(*t/λ*)^*k*^].
- *Gompertz* : *f* (*t*) ∝ *e*^*t/σ*^ exp[*c* (1 − *e*^*t/σ*^)].
- *Power law* : *f* (*t*) ∝ *t*^−*a*^.
- *Power law* + *cutoff* : *f* (*t*) ∝ *t*^−*a*^*e*^−*t/τ*^.
- *Stretched exponential* : *f* (*t*) ∝ exp[−(*t/τ*)^*β*^].

All candidate distributions were fitted using the same observation model, conditioned on the analysis window and incorporating a uniform contaminant with weight 0.02 (Eq. (5)). Rescoring these candidates under the numerically exact first-passage density, obtained by solving the Fokker-Planck equation with an absorbing boundary (Crank-Nicolson, d*x* = d*t* = 0.02), leaves all five inferior to the variable-start model for 98 to 100% of readers (*n* = 100, with median differences of 0.097 to 0.163 nats per read). The parameters were those selected under the approximation, so refitting could only improve them and the comparison is therefore conservative. Each candidate was scored on three quantities, computed on reads held out from its own estimation:

- The **held-out log-likelihood per read** is the expected log-probability the model assigns to a read it has not seen. In practice, it is estimated as the mean of log *f*_obs_(*t*) over the reads withheld from estimation, with *f*_obs_ the window-conditioned density of Eq. (5) evaluated at the parameters fitted to the other half.
- The **Kolmogorov-Smirnov distance** is the largest difference between the fitted and empirical cumulative distributions, *max* |*F*_model_ − *F*_*n*_|. It focus on the single worst region and is insensitive to how well the remainder fits.
- The **Wasserstein-1 distance** integrates the same discrepancy, ∫|*F*_model_ − *F*_*n*_ |d*t*, rather than taking its maximum like the Kolmogorov-Smirnov distance. It can be read as the average distance probability mass would have to move to turn one distribution into the other. Unlike the other two it is expressed in seconds, and is therefore the only one interpretable in the units of the data.

A model may rank differently on each, but when it does the disagreement is informative, indicating that candidates fail not only in different amounts, but also in different ways. Reporting a single one would hide that.The same models were compared at the three levels introduced in the main text, each of which tests something the others cannot.

In the **pooled** view (Table S1, Fig. S4), every candidate is fitted to all reads as though they came from a single reader. Its purpose is to rule out the objection that the result depends on fitting readers individually.

In the **individual** view (Table S2, Fig. S5), each candidate is fitted to each reader separately, and the fits are summarised in two ways. *Win* is the percentage of readers for whom the variable-start model possess the higher held-out log-likelihood in a pairwise contest with the candidate, and *Best* the percentage in which the candidate obtains the highest log-likelihood of all fourteen.

In the **mixture** view (Table S3, Fig. S6), the same per-reader fits are averaged with weights proportional to each reader’s number of reads. Candidates are compared through the difference in held-out log-likelihood computed within each reader rather than through scores pooled across readers.

Inference in the individual and mixture views uses the Wilcoxon signed-rank test on paired per-reader differences, Holm-corrected across candidates within each measure and window, with the matchedpairs rank-biserial correlation as the effect size. With 2820 readers, even negligible differences reach significance, so the effect size is the more informative quantity. The pooled view admits no such test, since each candidate yields a single fit and there is nothing to pair.

**Figure S4:**
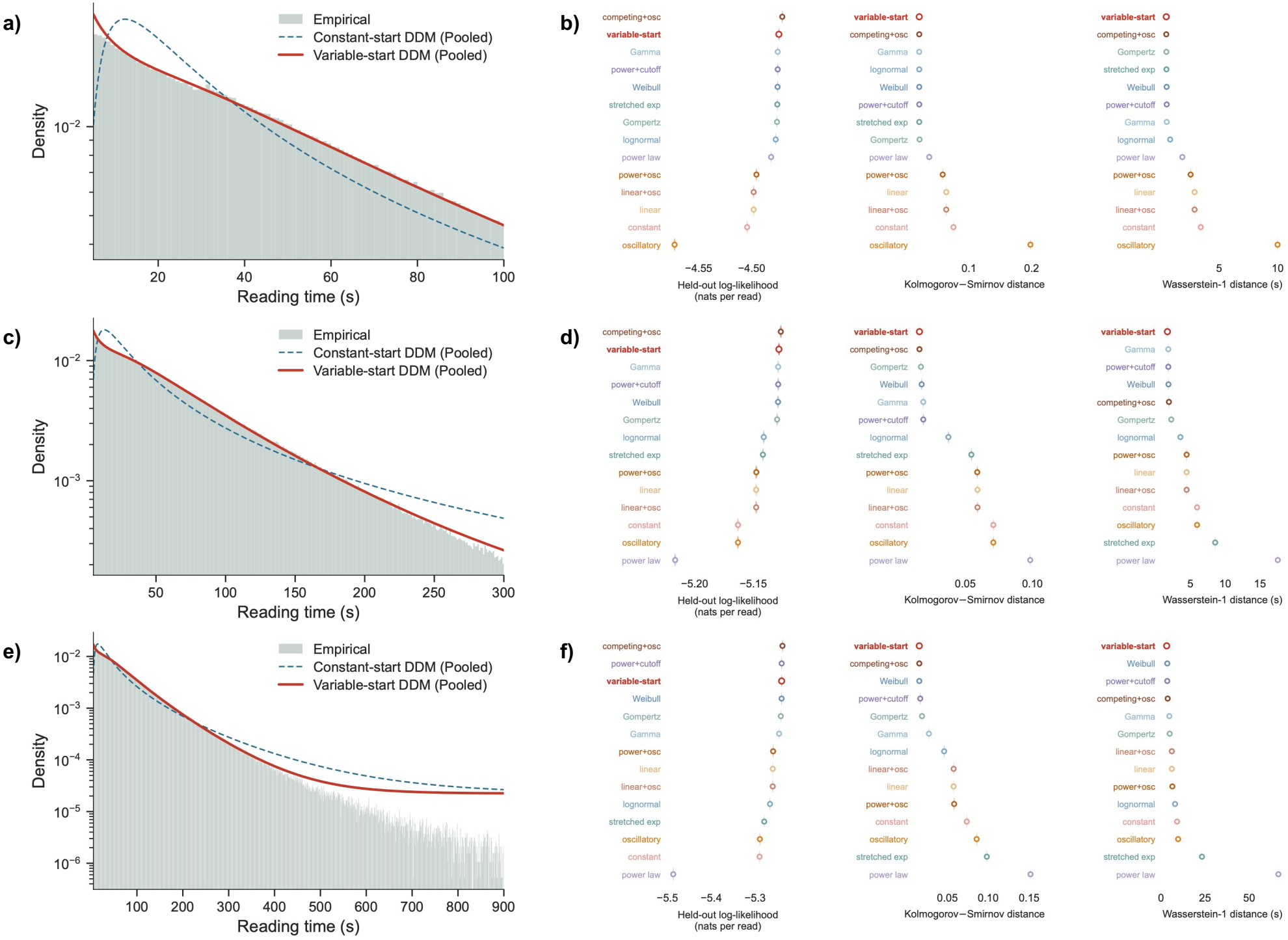
Pooled model comparison. Each candidate density was fitted to all article visits as though they came from a single reader, and scored on held-out reads. Rows of panels correspond to analysis windows of *t*_max_ = 100 s (**a, b**), 300 s (**c, d**) and 900 s (**e, f**), with all densities being conditioned on [5, *t*_max_] and include a uniform contaminant of weight 0.02. **a, c, e**, Pooled reading-time distribution (green) with the constant-start (dashed blue) and variable-start (solid red) drift-diffusion models (DDMs), which differ only in whether the starting point is fixed at zero or drawn afresh on each visit from *U*(0, *z*_frac_*θ*). Vertical axes are logarithmic. **b, d, f**, All fourteen candidates ranked on three measures: held-out log-likelihood per read (higher is better), Kolmogorov-Smirnov distance (the largest discrepancy between fitted and empirical cumulative distributions) and Wasserstein-1 distance (the same discrepancy integrated over the window, in seconds), for both distances lower is better. Candidates are ordered by each measure independently, drift-diffusion models are shown in warm colours and descriptive families in cool colours, with the variable-start model in bold. Single fits admit no per-reader pairing, so no statistical test is shown.

**Table S1:** Pooled model comparison. Each candidate fitted to all reads as though they came from a single reader, and scored on held-out reads. Values are absolute, with one fit per candidate and no per-reader pairing, so no paired test applies. Higher is better for the log-likelihood, lower for both distances. Bold marks the best value in each column and rows are ordered by the log-likelihood at *t*_max_ = 300 s.

|  |  | *t*<sub>max</sub> = 100 s |  |  | *t*<sub>max</sub> = 300 s |  |  | *t*<sub>max</sub> = 900 s |  |  |
|---|---|---|---|---|---|---|---|---|---|---|
| Candidate | Family | ll | KS | *W*<sub>1</sub> | ll | KS | *W*<sub>1</sub> | ll | KS | *W*<sub>1</sub> |
| Competing + oscillatory | DDM | **-4.4713** | **0.0206** | 0.50 | **-5.1266** | **0.0158** | 1.90 | **-5.2144** | **0.0156** | 3.80 |
| *Variable-start* | DDM | -4.4745 | **0.0206** | **0.50** | -5.1282 | **0.0158** | **1.69** | -5.2172 | **0.0156** | **3.21** |
| Gamma | Description | -4.4755 | **0.0206** | 0.54 | -5.1288 | 0.0187 | 1.82 | -5.2261 | 0.0273 | 4.72 |
| Power law + cutoff | Description | -4.4755 | **0.0206** | 0.54 | -5.1288 | 0.0187 | 1.82 | -5.2166 | 0.0167 | 3.54 |
| Weibull | Description | -4.4757 | **0.0206** | 0.53 | -5.1289 | 0.0175 | 1.84 | -5.2168 | **0.0156** | 3.54 |
| Gompertz | Description | -4.4763 | 0.0210 | 0.51 | -5.1297 | 0.0169 | 2.24 | -5.2165 | 0.0189 | 4.93 |
| Log-normal | Description | -4.4775 | **0.0206** | 0.83 | -5.1411 | 0.0374 | 3.55 | -5.2462 | 0.0460 | 7.96 |
| Stretched exponential | Description | -4.4759 | **0.0206** | 0.51 | -5.1418 | 0.0545 | 8.56 | -5.2637 | 0.0982 | 23.22 |
| Power + oscillatory | DDM | -4.4956 | 0.0580 | 2.57 | -5.1475 | 0.0588 | 4.45 | -5.2346 | 0.0582 | 6.44 |
| Linear drift | DDM | -4.4982 | 0.0637 | 2.91 | -5.1476 | 0.0590 | 4.45 | -5.2347 | 0.0576 | 6.14 |
| Linear + oscillatory | DDM | -4.4982 | 0.0637 | 2.91 | -5.1476 | 0.0590 | 4.45 | -5.2347 | 0.0576 | 6.14 |
| Constant-start | DDM | -4.5044 | 0.0752 | 3.44 | -5.1631 | 0.0708 | 5.95 | -5.2693 | 0.0736 | 9.11 |
| Oscillatory drift | DDM | -4.5728 | 0.1980 | 9.98 | -5.1631 | 0.0708 | 5.95 | -5.2682 | 0.0857 | 9.72 |
| Power law | Description | -4.4818 | 0.0365 | 1.87 | -5.2167 | 0.0983 | 17.56 | -5.4675 | 0.1516 | 66.56 |

For the pooled view and for *t*_max_ = 300 s, the competing + oscillatory drift-diffusion model obtains the highest pooled held-out log-likelihood, the variable-start model the lowest Wasserstein-1 distance, and the two possess the same value on the Kolmogorov-Smirnov distance (Table S1). No candidate dominates on all three measures, and the ordering is not stable across windows, but it should be noted that the variable-start model is at or near the best on both distances at every window. On the other hand, observe that this is the weakest of the three views, given that it was fitted to the aggregate and additional drift parameters can potentially absorb variation that arises between readers rather than within them.

**Table S2:** Individual model comparison. *Win* is the percentage of readers for whom the variable-start model obtains the higher held-out log-likelihood in a pairwise comparison with a given candidate. *Best* is the percentage for whom the said candidate obtains the highest log-likelihood of all fourteen models. The two count different events and neither follows from the other. The variable-start model has no win rate against itself. Readers for whom two or more candidates tie to within floating-point precision contribute a fraction to each in the *Best* column. Bold marks the highest share in each column and rows are ordered by *Best* at *t*_max_ = 300 s.

|  |  | *t*<sub>max</sub> = 100 s |  | *t*<sub>max</sub> = 300 s |  | *t*<sub>max</sub> = 900 s |  |
|---|---|---|---|---|---|---|---|
| Candidate | Family | Win (%) | Best (%) | Win (%) | Best (%) | Win (%) | Best (%) |
| Competing + oscillatory | DDM | 67 | **20.5** | 67 | **20.1** | 67 | 20.0 |
| *Variable-start* | DDM | — | 12.8 | — | 19.9 | — | **22.2** |
| Gompertz | Description | 62 | 16.2 | 65 | 15.6 | 68 | 13.0 |
| Power law + cutoff | Description | 47 | 4.2 | 49 | 9.9 | 49 | 10.0 |
| Gamma | Description | 47 | 4.5 | 49 | 8.4 | 49 | 11.5 |
| Weibull | Description | 49 | 6.1 | 48 | 7.7 | 48 | 7.7 |
| Stretched exponential | Description | 59 | 7.6 | 55 | 7.7 | 54 | 7.5 |
| Log-normal | Description | 63 | 7.2 | 88 | 4.8 | 96 | 2.2 |
| Power + oscillatory | DDM | 79 | 7.6 | 87 | 1.9 | 86 | 2.2 |
| Linear drift | DDM | 90 | 0.9 | 88 | 1.7 | 90 | 1.6 |
| Linear + oscillatory | DDM | 91 | 1.1 | 88 | 1.5 | 88 | 1.6 |
| Power law | Description | 75 | 9.8 | 99 | 0.4 | 100 | 0.1 |
| Constant-start | DDM | 93 | 0.7 | 97 | 0.3 | 98 | 0.1 |
| Oscillatory drift | DDM | 93 | 1.0 | 97 | 0.2 | 98 | 0.3 |

**Figure S5:**
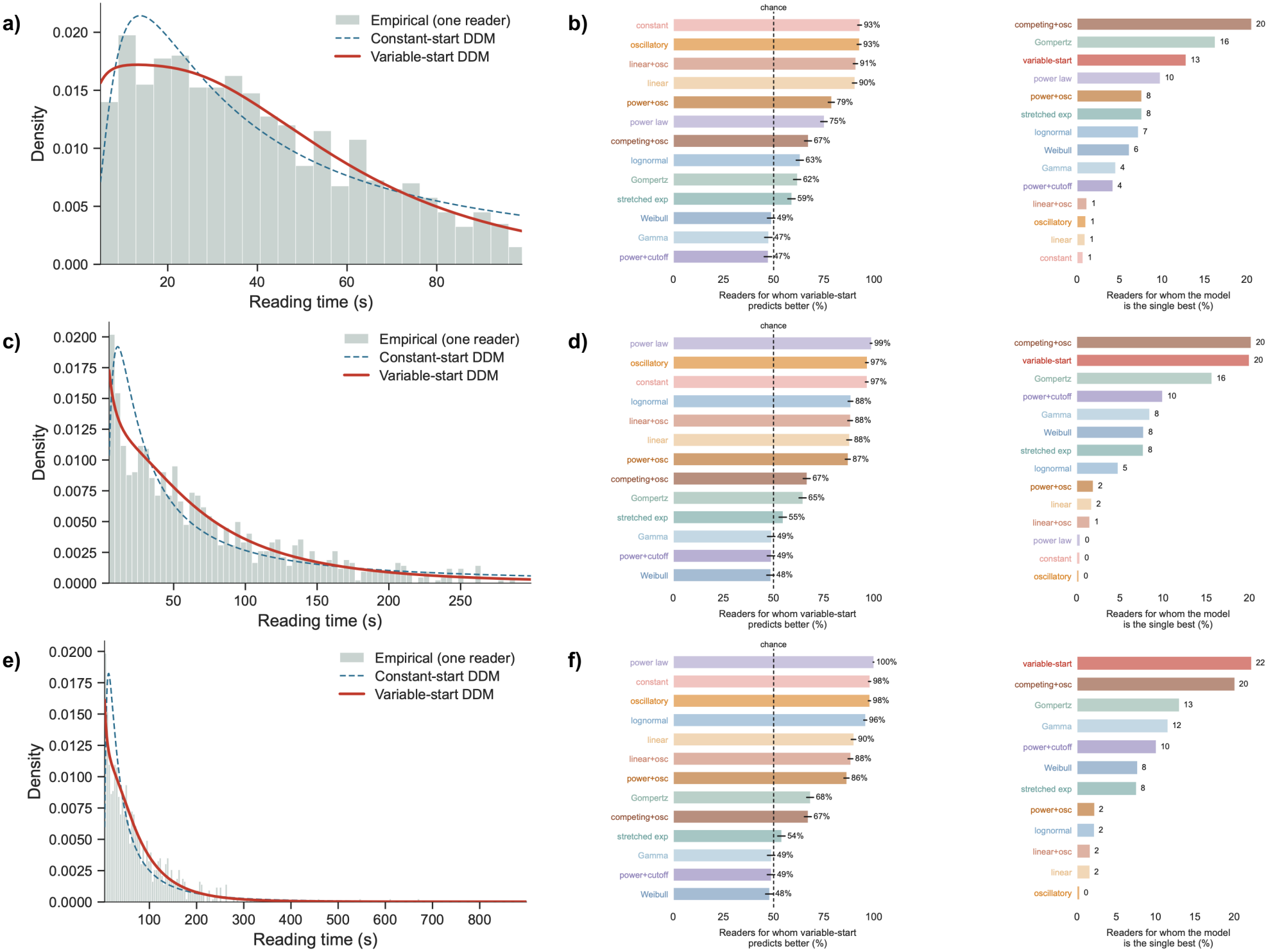
Individual model comparison. Each candidate was fitted to each reader separately, and scored on held-out reads. Panel rows correspond to *t*_max_ = 100 s (**a, b**), 300 s (**c, d**) and 900 s (**e, f**). **a, c, e**, Reading times of a single representative reader (green) with the constant-start (dashed blue) and variable-start (solid red) DDMs fitted to that reader’s own reads, with the reader selected at the median of fit among those in the top decile by read count, so selection was on sample size rather than on goodness of fit. Vertical axes are linear. **b, d, f**, Left, percentage of readers for whom the variable-start model attains the higher held-out log-likelihood in a pairwise contest with each candidate. The dashed line marks chance (50%). Right, percentage of readers for whom each candidate obtains the highest held-out log-likelihood of all fourteen. Readers for whom two or more candidates tie contribute a fraction to each. *n* = 2820 readers.

For the individual view, by counting how often each candidate is the single best of the fourteen, one observes the competing + oscillatory and variable-start models are nearly tied at *t*_max_ = 300 s (20.1% and 19.9% of readers respectively), with the Gompertz distribution third at 15.6%. The ordering between the two leading models depends on the window, with the competing + oscillatory model leading at 100 and 300 s and the variable-start model at 900 s (22.2% against 20.0%). That these two are close on this metric while the variable-start model wins 67% of their pairwise comparisons illustrates the distinction between the two statistics. Nevertheless, no candidate wins a majority of its pairwise contests against the variable-start model. In fact, the closest are the Gamma, Weibull and power law with exponential cutoff, which are indistinguishable from chance at 47 to 49%.

For the mixture view, by averaging the per-reader fits, the variable-start model obtains the best value on *all three* measures at *t*_max_ = 300 and 900 s (Table S3). It exceeds every other drift-diffusion model, including the competing + oscillatory one that leads the pooled comparison (median held-out log-likelihood −5.0647 against −5.0689 at *t*_max_ = 300 s, *p* < 0.001 on all three measures). The contrast between the pooled and mixture views suggests that the pooled fit was possibly absorbing betweenreader heterogeneity (once each reader is fitted separately, the additional drift parameters have nothing left to absorb). It also shows that the per-reader fits reproduce the aggregate distribution without any population-level parameter. Furthermore, a Gamma, a Weibull and a power law with exponential cutoff do not differ from the variable-start model on the log-likelihood or the Kolmogorov-Smirnov distance at *t*_max_ = 300 or 900 s, although all three are distinguishable on the Wasserstein-1 distance at 300 s. At *t*_max_ = 100 s the same three obtain nominally higher log-likelihoods and the difference reaches significance, while remaining indistinguishable on both distances. However, we read this shortest window as the least informative rather than as a counterexample, given that truncation at 100 s places a substantial part of many readers’ fitted densities outside the observed range.

**Figure S6:**
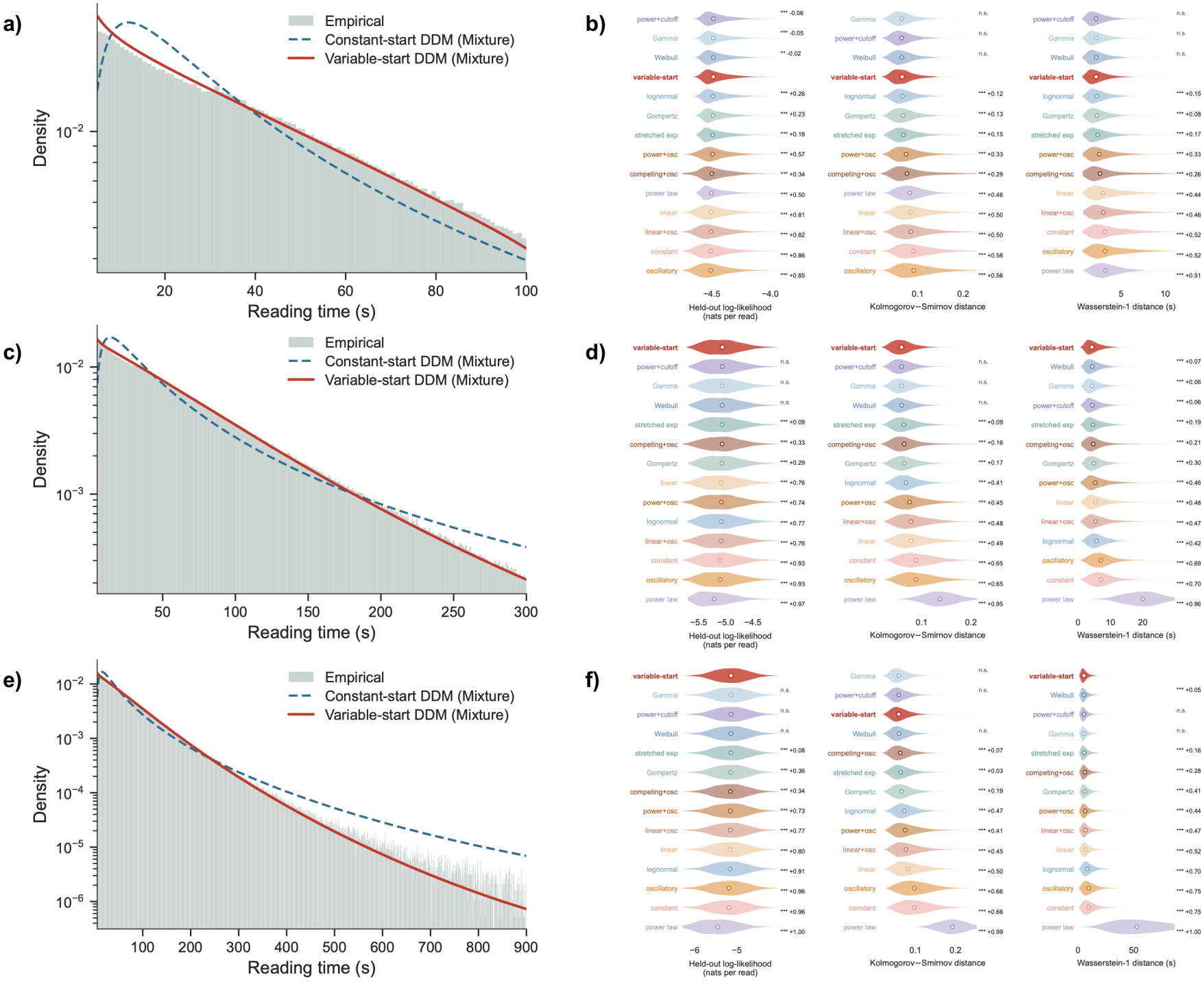
Mixture model comparison. Each candidate fitted to each reader separately and the resulting densities averaged with weights proportional to each reader’s number of reads. Panel rows correspond to *t*_max_ = 100 s (**a, b**), 300 s (**c, d**) and 900 s (**e, f**). **a, c, e**, Pooled reading times (green) with the read-weighted average of the individually fitted constant-start (dashed blue) and variablestart (solid red) densities. Vertical axes are logarithmic. **b, d, f**, Distribution across readers of each measure, for all fourteen candidates: held-out log-likelihood per read, Kolmogorov-Smirnov distance and Wasserstein-1 distance. Violins show the distribution over readers, open circles the median. Numbers to the right of each violin give the matched-pairs rank-biserial correlation of that candidate against the variable-start model, with positive values favouring the variable-start model. Asterisks give the Wilcoxon signed-rank test on the paired per-reader differences, Holm-corrected across candidates within each measure and window (^***^*P* < 0.001; n.s., not significant). *n* = 2820 readers.

The conclusion we draw is that goodness of fit does not separate an accumulation account from a sufficiently flexible description at the resolution under consideration, which we regard as a constraint on the argument rather than a weakness of the model. In other words, a two-parameter Gamma density summarises the reading times it was estimated from and tells us nothing about the reader, whereas *θ* and *α* are claims about a reader that can be tested elsewhere in that reader’s behaviour.

**Table S3:**
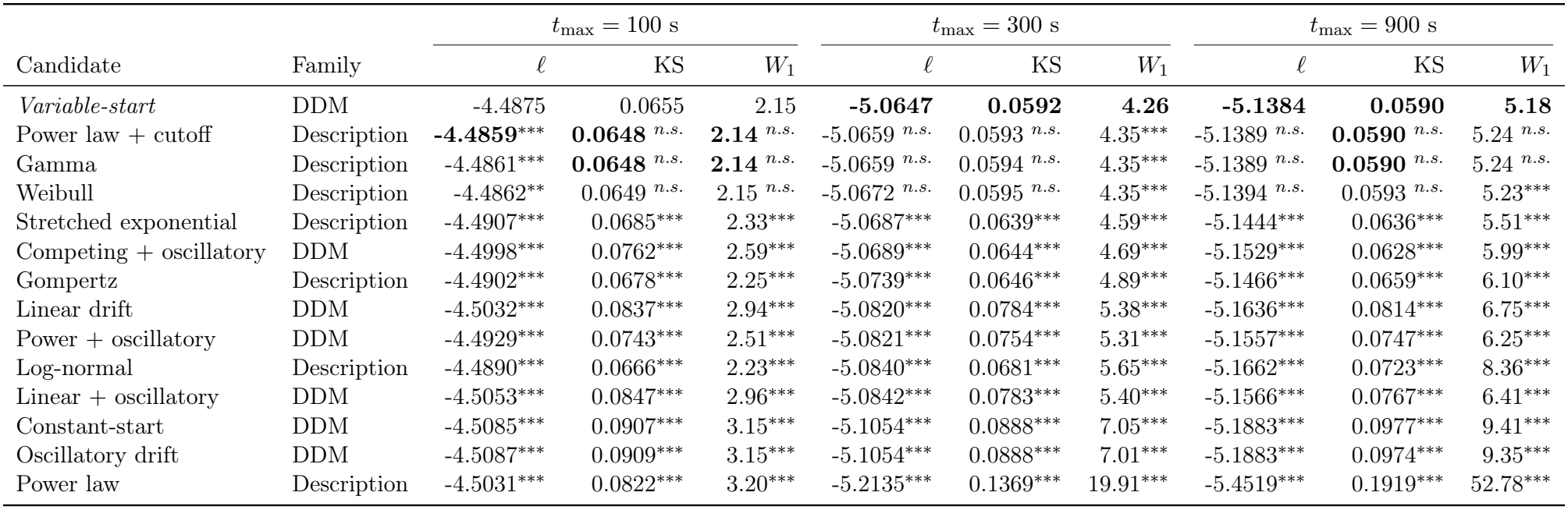
Mixture model comparison. Each candidate fitted to each reader separately and the resulting densities averaged with weights proportional to each reader’s number of reads. Entries are the median across readers of each measure, computed on held-out reads (*n* = 2820 readers). Higher is better for the log-likelihood, lower for both distances, bold marks the best value in each column and rows are ordered by the log-likelihood at *t*_max_ = 300 s. Markers give the paired comparison of that candidate with the variable-start model on that measure, by Wilcoxon signed-rank on the perreader differences, Holm-corrected across candidates within each measure and window: ^***^*p* < 0.001, ^**^*p* < 0.01, ^*^*p* < 0.05, n.s. otherwise.

## 4 Parameter identification

Despite the results presented in previous sections, a question remains unanswered. When the parameters are known, either by construction or by design, are we able to identify them correctly? Indeed, a fitted parameter is only interpretable if the model can recover it from data generated with that parameter. Furthermore, can the estimates obtained from part of the dataset be replicated using another part of the same dataset? The two questions address distinct problems related to parameter identification.

To address the first question, we generate synthetic readers using the fitted parameter values of individual real readers. Each synthetic reader is assigned the same number of reads as its real-data counterpart. These reads are drawn from the variable-start model using the corresponding fitted parameter values and are truncated to the analysis window. The resulting sample is then refitted using the same procedure employed throughout, with the analysis window being considered to ensure that generation and estimation use similar bounds. This allows us to assess whether the estimation procedure can recover the parameters used to generate the data. The recovered value is then compared with the value used to generate. Simulating at fitted values rather than over a parameter lattice was a deliberate choice. A lattice assess recovery at parameter combinations that no real reader occupies, while implicitly assigning them the same weight as combinations that are actually observed. By contrast, simulating at the fitted values of real readers keeps the assessment focused on parameter configurations represented in the data, making the recovery analysis directly relevant to the empirical population under study.

Patience and drift rate are recovered across their whole range (Fig. S7, with Spearman *ρ* = 0.972 and 0.899 respectively). The median recovered value differs from the true one by 1.4% for patience and 0.8% for the drift rate, and for half of readers the recovered value falls between −8% and +5% of the true one for patience, and between −7% and +6% for the drift rate. The assessment floor behaves differently, with its ordering properly recovered (*ρ* = 0.717) but not necessarily its value. Alongside the diagonal, Fig. S7c) shows two off-diagonal groups of readers, those with a substantial true floor recovered near zero and those with a near-zero true floor recovered at a substantial one, which together produce a ratio of recovered to true floor with an upper quartile above 300 against a median of 1.074.

**Figure S7:**
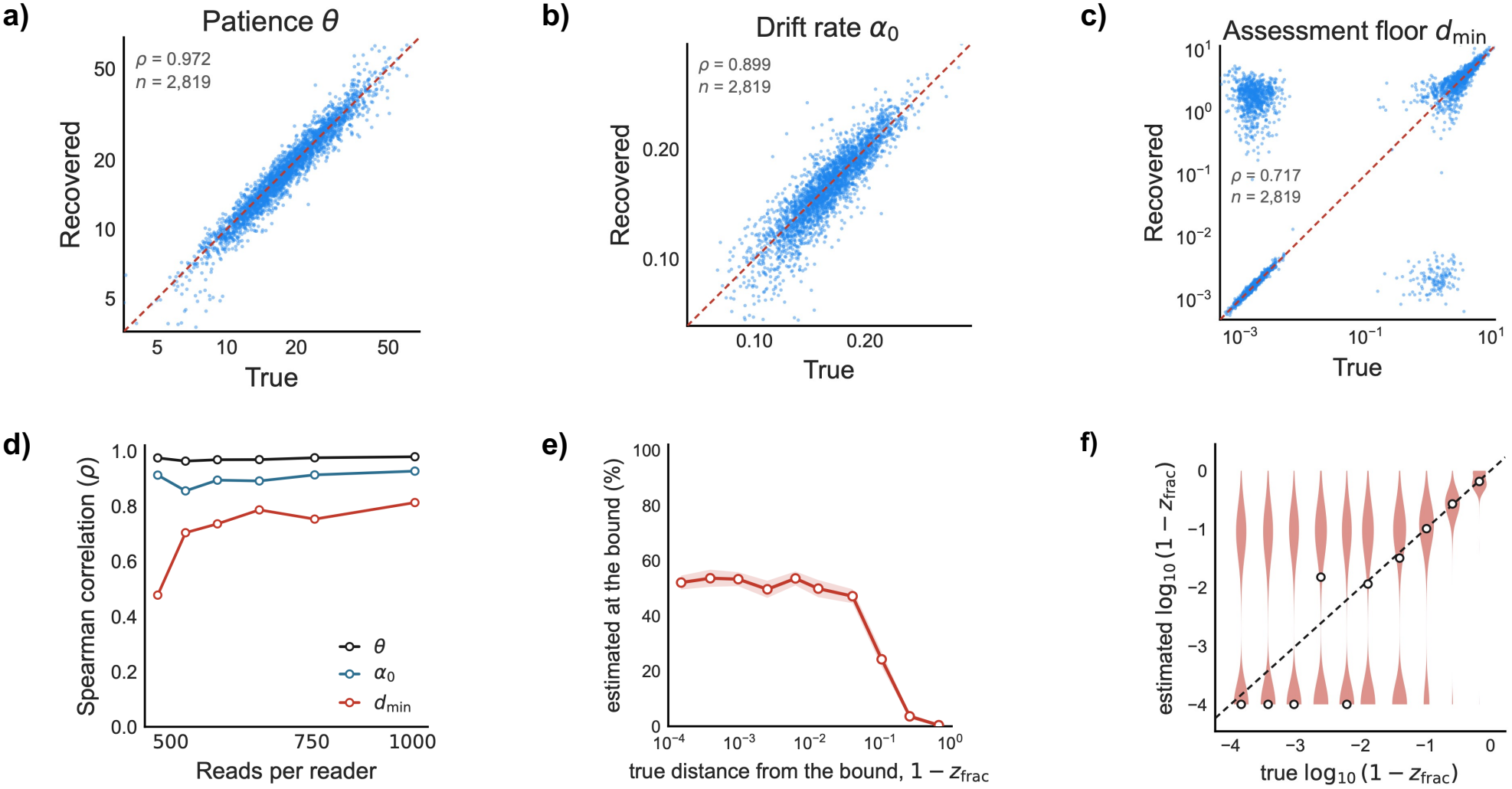
Parameter identification on synthetic readers. Each synthetic reader was generated at the fitted patience, drift rate and start fraction of a real reader, with the same number of reads that reader contributed, truncated to the [5, 300) s analysis window and refitted by the procedure used throughout. **a), b), c)**, Recovered against true values for patience *θ*, drift rate *α*_0_ and the derived assessment floor *d*_min_ = *θ*(1 −*z*_frac_), with the identity line dashed and the Spearman correlation inset. Axes are logarithmic. **d)**, Spearman correlation between true and recovered values within bins of read count, each coloured line representing a parameter. For panels **e)** and **f)**, synthetic readers were generated at different distances from the upper bound of the start fraction rather than at fitted values, since the fitted distribution is concentrated at the bound and leaves the intervening range almost empty. **e)**, Proportion of readers whose fitted start fraction exceeds 0.99, against the true distance from the bound, with 500 readers for each point. The shaded band is a 95% confidence interval on the proportion, from the normal approximation to the binomial. **f)**, The full distribution of recovered values at each true distance, as violins on logarithmic axes with the identity line dashed and the median marked. The recovered values form two separated groups rather than a continuum around the truth, a structure the proportion in **e)** summarises but cannot display.

Binning on read count separates the three parameters (Fig. S7d). Patience and drift rate are recovered almost equally well at every sample size, which indicates that their recovery is a property of the model rather than of how much data a reader provided. The assessment floor is not, rising from around 0.5 at the sparsest bin to around 0.8 at the densest, so its weaker recovery is partly a matter of sample size.Sample size does not account for the floor’s weaker recovery on its own. Even at the densest bin it is recovered less well than either of the other parameters, so what remains is a property of the quantity rather than of the data available to estimate it. (Fig.S7e) isolates that by asking how close to its bound the start fraction can lie and still be told apart from it. Unlike the other panels, Fig. S7e does not use the start fraction values fitted to real readers. Simulating from these values reproduces the sample’s own distribution of the start fraction, which is piled at the bound and almost empty between there and 0.1, which is the region the panel has to resolve. Synthetic readers were therefore generated at different prescribed distances from the bound, with 500 readers at each, drawn from the fitted patience and drift rate of real readers.

**Table S4:** Reading times generated at two start fractions spanning the plateau. For each pair of patience and drift rate, 100000 reads were generated at *z*_frac_ = 0.9999 and at 0.95, truncated to the [5, 300) s analysis window, and the two distributions compared by a two-sample Kolmogorov-Smirnov statistic. The two are indistinguishable across the range of patience the sample occupies (median 17.8, 90th percentile 30.6, 99th percentile 46.7) and separate only above it.

| *θ* | 0.1 |  |  | 1 |  |  | 5 |  |  | 10 |  |  | 20 |  |  | 40 |  |  | 80 |  |  |
|---|---|---|---|---|---|---|---|---|---|---|---|---|---|---|---|---|---|---|---|---|---|
| *α*<sub>0</sub> | .08 | .16 | .32 | .08 | .16 | .32 | .08 | .16 | .32 | .08 | .16 | .32 | .08 | .16 | .32 | .08 | .16 | .32 | .08 | .16 | .32 |
| KS | .002 | .002 | .002 | .003 | .002 | .002 | .002 | .002 | .002 | .002 | .002 | .001 | .005 | .004 | .004 | .015 | .012 | .008 | .047 | .030 | .017 |
| under 10 s, bound | 34.2 | 40.3 | 54.6 | 33.4 | 39.6 | 53.7 | 22.8 | 28.1 | 41.4 | 11.3 | 13.8 | 21.5 | 5.4 | 6.1 | 9.4 | 3.5 | 3.1 | 4.4 | 3.2 | 2.2 | 2.2 |
| under 10 s, *z*<sub>*frac*</sub> = 0.95 | 34.3 | 40.2 | 54.4 | 33.2 | 39.5 | 53.9 | 22.6 | 28.1 | 41.4 | 11.3 | 13.8 | 21.6 | 5.1 | 5.9 | 9.1 | 2.7 | 2.6 | 3.9 | 1.3 | 1.0 | 1.2 |

In practice, (Fig. S7e) plots the proportion of synthetic readers whose fitted start fraction exceeds 0.99, against the true distance from that bound. It is plotted against 1 −*z*_frac_ rather than against *d*_min_ deliberately because, since *d*_*min*_ = *θ*(1 −*z*_frac_), a falling detection rate would conflate the distribution of patience with the limits of identification.The rate is flat at about half across three orders of magnitude, from the bound itself to around 0.05, and only then falls, becoming negligible beyond around 0.3. Within that plateau the fitted value carries almost no information about the true one, since a reader at the bound and a reader at a distance of 0.05 from it are placed at the bound with similar frequency. Beyond it the rate decreases, so a fitted value far from the bound is informative where one close to it is not. Fig. S7f) shows the same comparison without reducing each reader to whether their fitted start fraction exceeds 0.99. At every true distance inside the plateau the estimated values fall into two separate groups, one around the bound and one well above it with little between, and the median sits around the bound until the true distance exceeds about 0.01. The detection rate in panel e) counts the readers above that threshold, which cuts through the lower group rather than around it. What the violins add is that the two groups are distinct rather than a spread around the truth, so a reader is placed firmly in one or the other and the true value does not decide which.

A parameter that cannot distinguish between values spanning three orders of magnitude is a serious limitation only if those values correspond to different behaviours, which they do not in our case, since generating reading times at the two ends of the plateau, *z*_frac_ = 0.9999 and 0.95, gives distributions that are barely distinguishable across the range of patience the sample occupies (Table S4). The Kolmogorov-Smirnov statistic between them is 0.002 at a patience of 10 and 0.005 at 20, and the share of reads ending within 10 s is also identical. The two separate only above a patience of about 40 (corresponding to the 98th percentile of the fitted values), and only 29 readers with a start fraction in the affected range (1% of the sample) lie above this threshold. Readers inside the plateau are therefore similar in the behaviour the model describes, whatever their fitted values. Patience and drift rate are identified, but the floor is identified in ordering (which is all the main text uses) and not in value near the bound. The conditional group is therefore described by the absence of a detectable floor, rather than by a small floor value. This is what the data support, rather than the claim that the floor is simply small.

Identification is not able to assess whether a reader’s fitted parameters are a stable property of that reader or an accident of which reads happened to be recorded. That is the second question, and it is not answered with simulations, but by comparing a reader’s data against itself. To do this, each reader’s reads inside the analysis window were divided randomly into two disjoint halves.The model was fitted independently to each half, and the two estimates compared across readers. No ground truth is available or needed, since agreement between halves is evidence that the estimate reflects the reader and disagreement that it reflects the sample.

Patience and drift rate replicate across halves (Fig. S8a and b, with Spearman correlations *ρ* = 0.887 and 0.780 across 2820 readers). The start fraction replicates less closely (Fig. S8d, with *ρ* = 0.646), and the derived assessment floor more so than the parameter it is derived from (Fig. S8e, with *ρ* = 0.793). The floor is displayed separately for that reason, since its reliability is not a function of the reliabilities of patience and the start fraction and has to be measured directly.Fig. S8e) has the same structure as its counterpart in the identification figure (Fig.S7c). In a similar fashion, alongside the diagonal sit two off-diagonal groups, readers whose floor is near zero in one half and substantial in the other, and the reverse. These are the readers for whom the two halves of their own reading support different conclusions about whether a floor is present.

**Figure S8:**
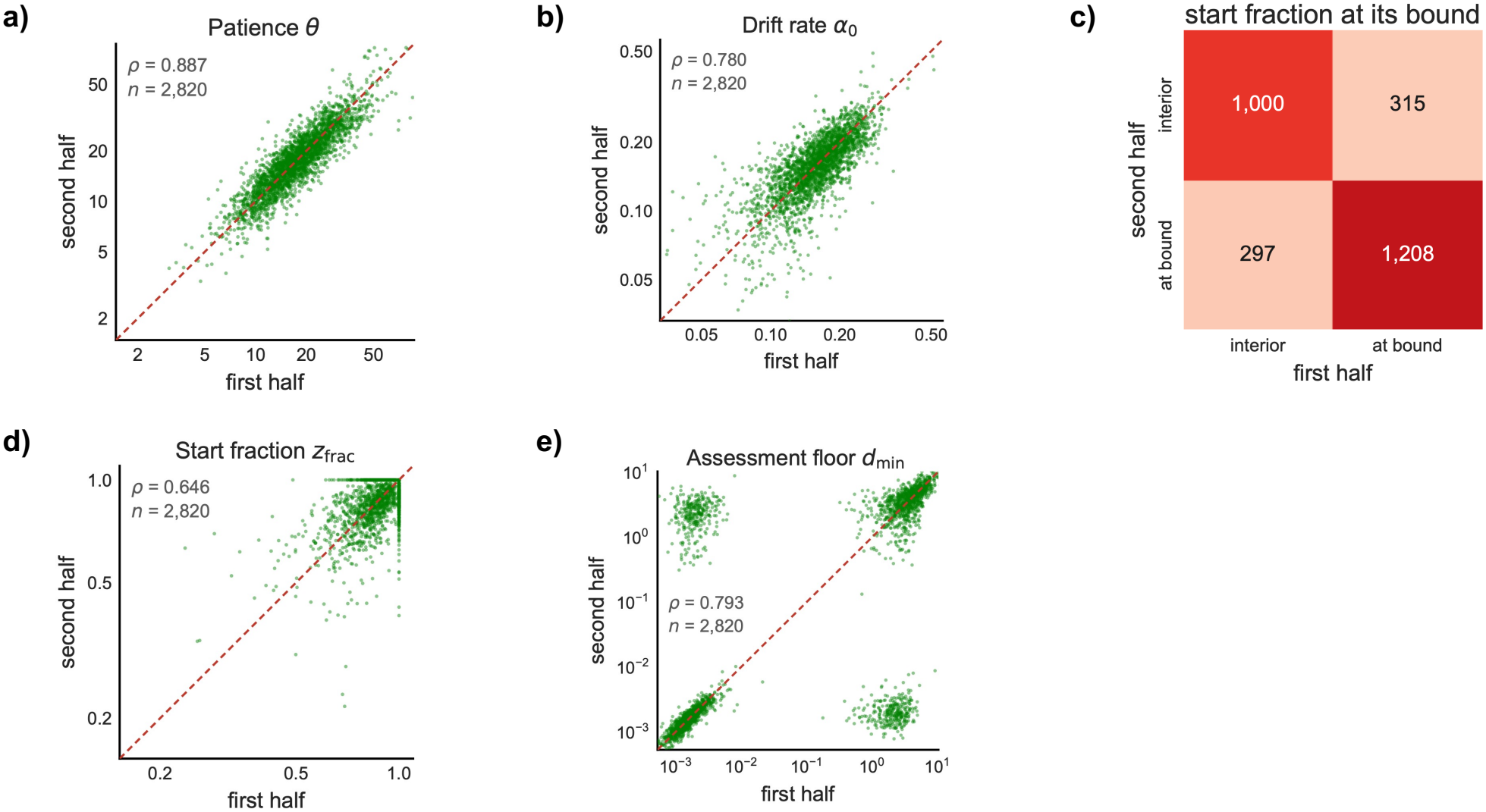
Stability of the fitted parameters across halves of a reader’s own reads. Each reader’s reads inside the [5, 300) s analysis window were divided at random into two disjoint halves and the model fitted independently to each. **a), b), d), e)**, Estimates from the first half against those from the second for patience *θ*, drift rate *α*_0_, the start fraction *z*_frac_ and the derived assessment floor *d*_min_ = *θ*(1− *z*_frac_), with the identity line dashed and the Spearman correlation inset. Axes are logarithmic. **c)**, Readers counted by whether their start fraction is bigger than 0.99 (at bound) in the first half against whether it is bigger in the second half. The off-diagonals represent cells in which the readers’ halves support opposite conclusions regarding the start fraction, while the diagonals represent cells supporting similar ones.

With this in mind, Fig.S8c) counts those readers directly, setting whether a reader’s start fraction exceeds 0.99 in one half against whether it does in the other. The agreement rate is not interpretable on its own, since with about half of readers above that value two independent draws weighted like the marginals would agree roughly half the time by construction. Cohen’s *κ* subtracts that baseline. The two halves agree for 78.3% of readers against a chance level of 50.3%, giving *κ* = 0.563, and 79.3% of readers above 0.99 in one half are above it in the other. Patience is a property of the reader rather than of the particular reads recorded, and so, more weakly, are the drift rate and the assessment floor. The start fraction replicates least well of the four, and the readers it fails on are those in the off-diagonal groups of Fig. S8e), for whom one of the two halves places the start fraction at its bound and the other does not.

Importantly, these readers are not behaving inconsistently, since the Kolmogorov-Smirnov statistic between their two halves of reads has a median of 0.058 for those whose halves disagree, against 0.047 for readers whose halves agree. The difference lies then in the estimate, and not the behaviour, which is the plateau the preceding figure reached by simulation, obtained here without one.

One number from the preceding two figures remains unexplained. On simulated readers generated at the bound, the estimator places only about half above 0.99. The same pattern should make two halves of a real reader’s data disagree about the start fraction just as often, and instead they agree for 79.3% of readers. Real reads drive the start fraction to its bound more consistently than reads simulated from the fitted parameters do, which implies the data contain something the model does not generate.

**Figure S9:**
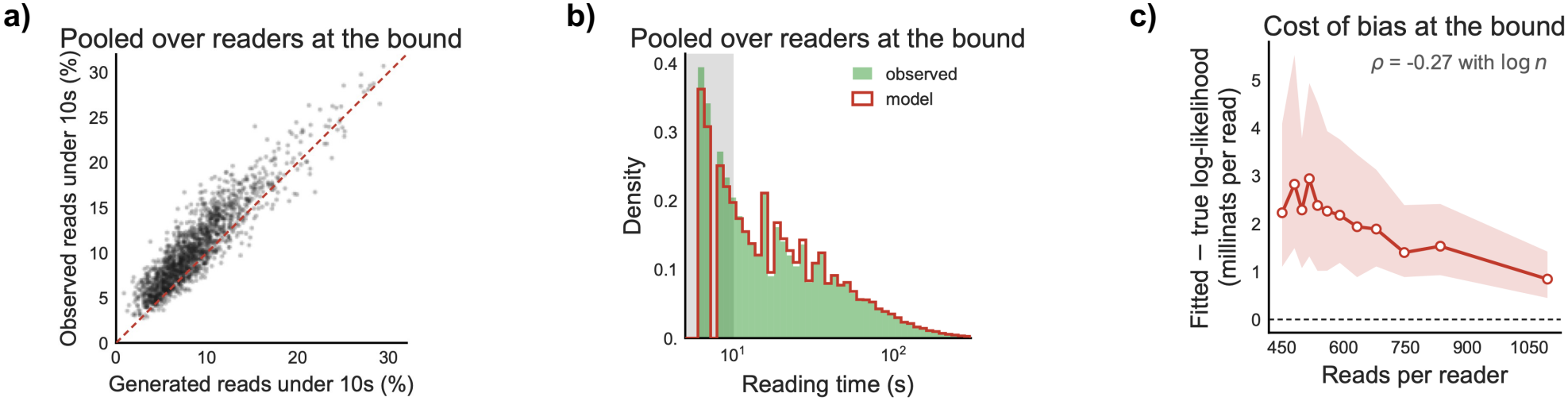
Comparison of observed reading times with reading times generated by fitted models. For each reader whose start fraction exceeds 0.99, a synthetic sample of the same size was generated at that reader’s own fitted parameters and truncated at the [5, 300) s analysis window. **a)** Observed versus generated proportion of reads under 10 s (one point per reader and identity line dashed). Points above the identity line are readers whose own readings possess more short visits than their synthetic counterparts. **b)** The two reading time distributions pooled across readers, with the 5 s to 10 s interval shaded. The reading time axis is logarithmic. **c)** The amount by which the fitted optimum exceeds the likelihood at the true parameters (in millinats per read) as a function of the number of reads the reader contributed, for synthetic readers generated at the bound. The line is the median within bins of read counts and the band the interquantile range. A positive value means that the fraction below the truth fits the generated reads better than the value they were generated at.

For each reader whose fitted start fraction exceeds 0.99, a synthetic sample of the same size was generated at that reader’s own fitted values, and the real and synthetic samples compared, first by the proportion of reads ending within 10 s and then by the pooled distribution of reading times. Because each synthetic reader has the same parameters and the same number of reads as its real counterpart, any difference in shape can then be attributed to the model rather than to differences between readers. The model under-generates the shortest reads (Fig. S9a). Across readers at the bound, 10.4% of observed reads end within 10 s against 8.7% of simulated reads, and the model generates fewer of them for 82% of readers. The discrepancy is confined to that range (Fig. S9b). Over all reads, the Kolmogorov-Smirnov statistic between the observed and simulated distributions is 0.018, but it decreases to 0.007 when restricted to reads of 10 s and above and increases to 0.093 when restricted to reads below 10 s. The model reproduces a reader’s reading times everywhere except at the left edge of the distribution.This discrepancy has a measurable cost (Fig. S9c). For simulated readers whose true start fraction is at the bound, a start fraction below it fits better, by a median of 1.8 millinats per read, which is about 1 nat over a typical reader’s reads. The margin narrows as readers contribute more reads (rank correlation −0.27), so it is a finite-sample effect, but it persists across the sample sizes available here.

A reader who produces more short reads than the model generates at their own parameters provides evidence for an assessment floor close to zero. That evidence is present in both halves of their reading, whose reads differ no more than any other reader’s (median Kolmogorov-Smirnov statistic between halves of 0.058, against 0.047 for readers whose fits agree). Each half is therefore fitted at the bound, and the two agree more often than simulation would predict.The misspecification is localised, and it does not weaken the interpretation given in the main text. In fact, a model that generates too few short reads makes the bound harder to reach, so readers placed there have more evidence behind them than the fit implies. What it affects is the precision of an individual reader’s fraction being exactly at the defined bound, which the main text never reports. The main results rest on the ordering of the floor, which replicates across halves at *ρ* = 0.793, and on a division between groups that a simple threshold on the start fraction recovers almost exactly, as we will see in the next section.

## 5 Clustering selection and robustness

The main text classifies readers into two groups and claims that this division predicts behaviour. However, an important question is whether alternative, equally reasonable methodological choices might have produced a different partitioning of the readers’ behavioural patterns. This question can be broken down into two smaller, more specific ones. First, how the number of components was selected, and second, how sensitive the resulting partition is to variations in the choices made during its construction.

**Figure S10:**
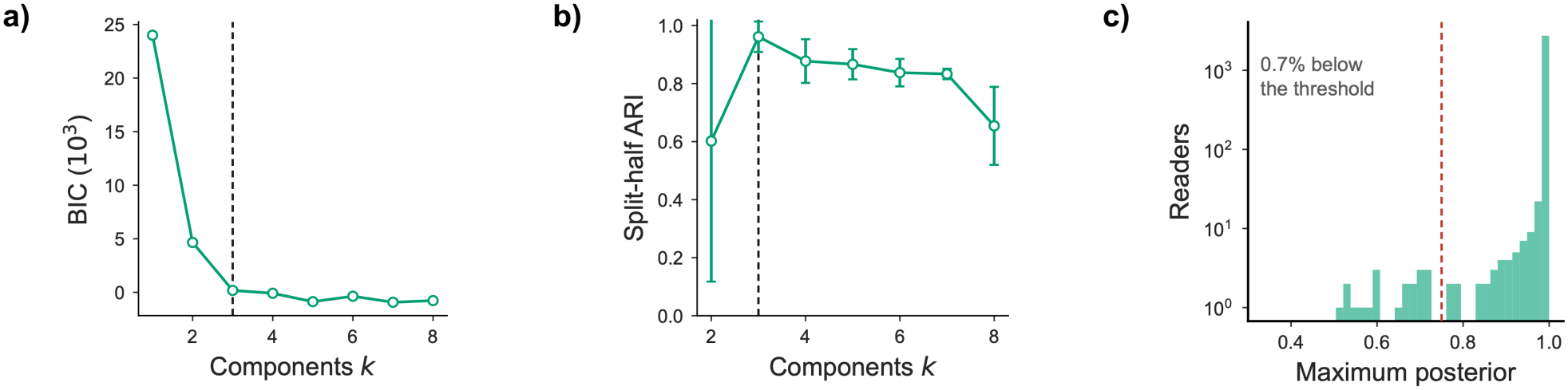
Selection of the number of mixture components. Gaussian mixtures with full covariance matrices were fitted for *k* = 1 to 8 on the log-transformed patience and drift rate together with the untransformed start fraction. The dashed vertical line in **a** and **b** marks the selected value, *k* = 3. **a)** Bayesian information criterion (BIC) as a function of *k*. BIC decreases over the first few values and then varies without a clear trend, reaching its lowest value at *k* = 7. **b)** Split-half reproducibility against *k*, measured as the adjusted Rand index (ARI) between the labellings assigned to the full sample by two mixtures, each fitted to one half of a random split of readers. Points are the mean and error bars the standard deviation across ten splits. The index peaks at *k* = 3. **c)** Distribution across readers of the maximum posterior probability of component membership, for the selected mixture. The dashed lines mark the assignment threshold of 0.75 and the readers below it were left unassigned rather than allocated to the closest component.

To obtain said partition, Gaussian mixtures with full covariance matrices were fitted for *k* = 1 to 8 on the log-transformed patience and drift rate together with the untransformed start fraction. The Bayesian information criterion (BIC) falls steeply from *k* = 1 to *k* = 3 and then varies without a clear trend, reaching its lowest value at *k* = 7, Fig. S10 a). The preference for seven components reflects the parameter distributions not being Gaussian (further components capture their shape rather than identifying further groups of readers). So, although the selection by BIC is the conventional choice, it is not available here.

The number of components was therefore selected by split-half reproducibility, Fig. S10 b). A mixture was fitted to each half of a random split of readers, both half-sample models were used to label the full sample, and the adjusted Rand index (ARI) between the two labelings was computed. The process was repeated over ten splits, and the error bars are the standard deviation across them. The index peaks at 0.96 for *k* = 3, above the values at other numbers of components, and this number was therefore selected. Note that, because both half-sample models label the full sample, the ARI compares two labelings of the same readers rather than of disjoint sets, and only the comparison across *k* matters for the selection. Moreover, BIC and reproducibility answer different questions. A mixture with seven components describes the parameter distributions more closely than one with three, which is what BIC represents, while assigning readers to components in a way that does not survive refitting on half the sample. Where the objective is a division of readers rather than a description of a density, reproducibility is the property that matters. The mixture returns, for each reader, a probability of belonging to each component. Readers whose largest maximum posterior probability fell below 0.75 were left unassigned rather than forced to the nearer component, which applies to 0.7% of readers. The mixture was fitted to patience, the drift rate and the start fraction, but the division it returns separates readers by only one quantity. Hartigan’s dip test, applied to the assessment floor and to the two fitted scale parameters across all readers with fitted estimates, locates the departure from unimodality, Fig. S11a), b) and c). The assessment floor (*D* = 0.151, *P* < 10^−3^) rejects unimodality, while patience (*D* = 0.005, *P* = 0.99) and the drift rate (*D* = 0.008, *P* = 0.29) do not.

**Figure S11:**
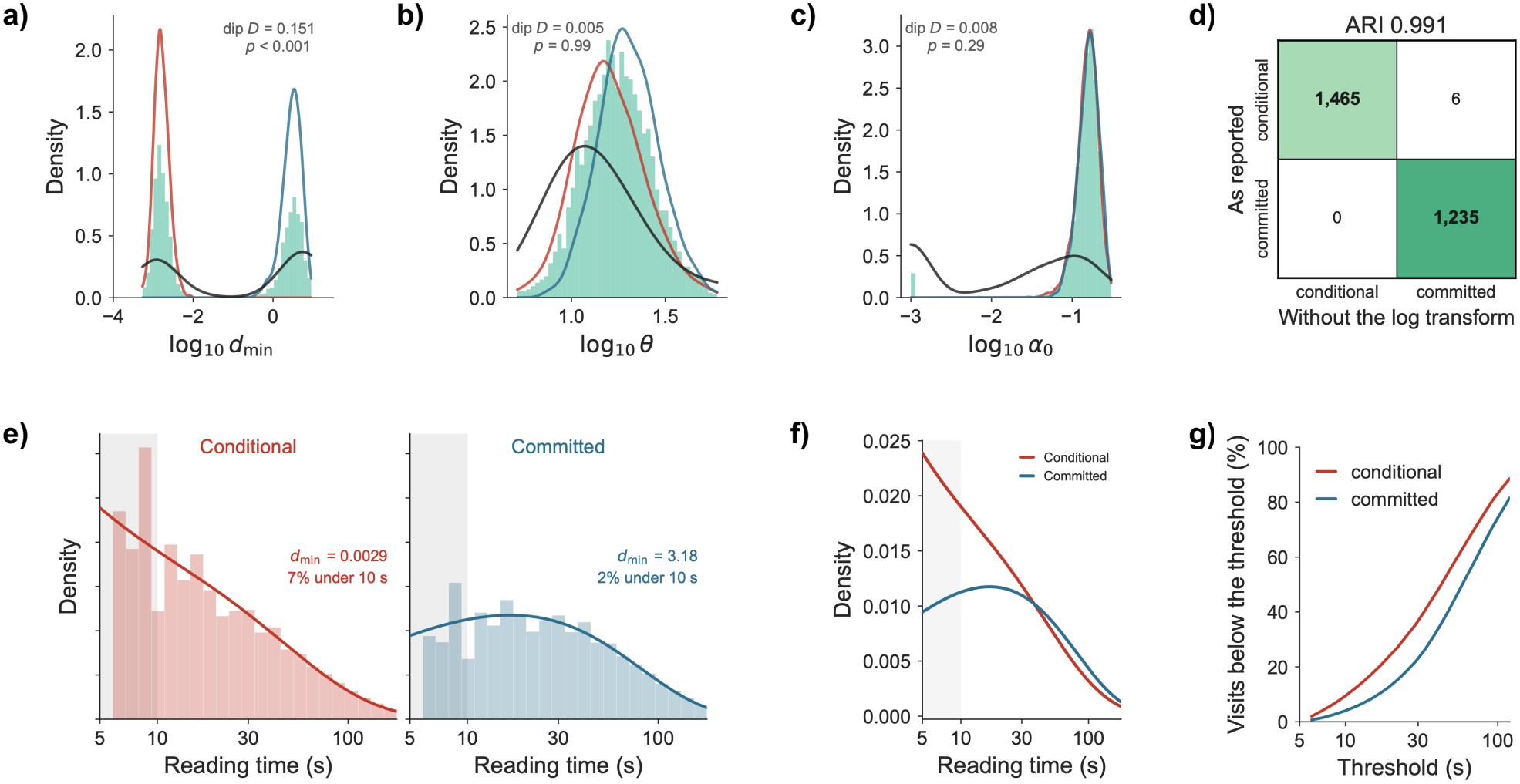
Robustness of the reader division. Throughout the figures, the conditional group is shown in red, the committed group in blue, and the readers of the excluded third component in black (the histograms in **a, b**, and **c** show all readers with fitted parameters, in green). **a), b), c)**, Distributions of the assessment floor (*d*_min_), patience (*θ*) and drift rate (*α*_0_) across all readers with fitted parameters, shown as a histogram with each group’s kernel density estimate drawn over it. Hartigan’s dip statistic *D* and its *P* value are inset. The assessment floor is bimodal (*D* = 0.151, *P* < 10^−3^) while patience (*D* = 0.005, *P* = 0.99) and the drift rate (*D* = 0.008, *P* = 0.29) are not. **d)** Agreement between the reported partition and one obtained by clustering the untransformed parameters, as counts of readers. Cells on the diagonal give the readers assigned in the same way by both and off-diagonal cells give those assigned differently. **e)** Reading times of each group pooled over all of its readers, with the read-weighted mean of that group’s readers’ own fitted densities drawn over the histogram (the shaded band marks the interval below 10 s). **f)** The same two densities of **e)** on one shared axis of the reading time. **g**, Share of a reader’s visits ending below each threshold, computed per reader and summarised by the median (bootstrap percentile intervals over readers reach a maximum width of 2.0 percentage points and are omitted for legibility).

Each panel draws the distribution for all readers with the three groups’ densities over it. In fact, the two clusters occupy distinct regions of the assessment floor and overlapping regions of patience and the drift rate, which encapsulates what the main text claims from the mixture, that the division is produced by the assessment floor. The excluded readers span both modes of the assessment floor, which is why the mixture could not place them on one side of the division. On patience they are indistinguishable from the two clusters. On the drift rate they are not, and a subset of them sit at values well below either cluster, which is why the dip statistic for the drift rate shifts from *P* = 0.29 across all fitted readers to *P* = 0.998 across the classified subset alone (a substantial change for 114 readers to produce, and one that helps distinguish them). A group distinguished by unusually slow evidence accumulation would be worth characterizing in its own right, but we do not do that here because, with only 114 readers, it is too small to support a description of a third reading strategy. Note that repeating this analysis on the classified subset rather than on all fitted readers leaves both statistics essentially unchanged (0.151 to 0.152 for the assessment floor, 0.005 for patience in both cases), so the departure from unimodality is a property of the sample and not a product of having excluded the readers the mixture could not place.

Patience and the drift rate were log-transformed before the analysis, so that equal ratios could correspond to equal distances. This is an assumption about the geometry a mixture with full covariances makes, and a reader may reasonably ask whether it produced the division rather than revealing it. Clustering the untransformed parameters answers the question and the same readers fall in the same groups, with 6 of the 2,706 moving from the conditional to the committed cluster (ARI = 0.991, Fig. S11d).

**Table S5:** Spearman rank correlations among the fitted parameters and the two behaviours, across the 2706 classified readers.

|  | *θ* | *d*<sub>min</sub> | navigation | abandonment |
|---|---|---|---|---|
| *θ* | X | +0.564 | −0.336 | −0.657 |
| *d*<sub>min</sub> | +0.564 | X | −0.176 | −0.817 |
| navigation | −0.336 | −0.176 | X | +0.074 |
| abandonment | −0.657 | −0.817 | +0.074 | X |

The main text illustrates the two groups with one representative reader from each, and whether those two are representative is a fair question. Pooling every retained read of every reader in each group settles this discussion, Fig. S11e). The conditional group’s distribution is highest at the shortest observable times and falls from there, with 7% of reads under 10 s, while the committed group’s rises to a later peak, with only 2% of the reads occurring below 10 s. These are the shapes the exemplar pair displays. Note that the curve in each panel of Fig. S11e) is the read-weighted mean of that group’s readers’ own fitted densities, not the density at the group’s median parameters. Drawn on one axis, Fig. S11f), the two densities separate below 10 s and tend to converge above about 30 s. The comparison is direct here rather than across two panels, and the shape of the difference is what matters. Patience sets the total evidence a decision requires, so a difference in it moves the whole distribution rather than only the left tail. The assessment floor sets the minimum, so a difference in it removes the fastest completions and leaves the rest close to where they were, since a visit that would have run a minute is barely affected by starting a little further from threshold. The difference between the two groups is therefore of the second kind, which is why their densities part at the left edge and not throughout.

Abandonment is reported as the share of visits ending within ten seconds. The threshold was defined in advance and not tuned, but a reader cannot establish this from this number alone, and a difference present only near this value would be a consequence of the choice itself rather than a property of the readers themselves. Fig. S11g) therefore gives the share of visits below every threshold from 6 to 120 s, computed per reader to match how the main text reports it and summarised by the median. The two curves remain separated across the range, so the division separates readers by how soon they leave an article rather than by a coincidence at a given cut-off.

The partial correlations reported in the main text are computed from the rank correlations below, with abandonment counted over all of a reader’s reads rather than only those inside the analysis window. Patience and the assessment floor correlate at 0.564, which is why partials are reported rather than the marginal correlations above. Since *d*_min_ = *θ*(1− *z*_frac_), a reader with a higher threshold tends also to have a higher floor, so either parameter will tend to correlate with a behaviour the other governs.

In conclusion, the division remains after removing the transformation, and does not depend on the particular readers used to illustrate it. It also appears in the pooled reads of every reader in each group and persists across all thresholds used to define abandonment. Taken together, these checks show that no single step or choice in the analysis is responsible for the clustering results we got. With that being said, these checks do not establish that the division is by itself correct, and no robustness check can. What supports the interpretation is the behaviour the division predicts, which is in the main text.

## 6 Independent testing in a different dataset

**Figure S12:**
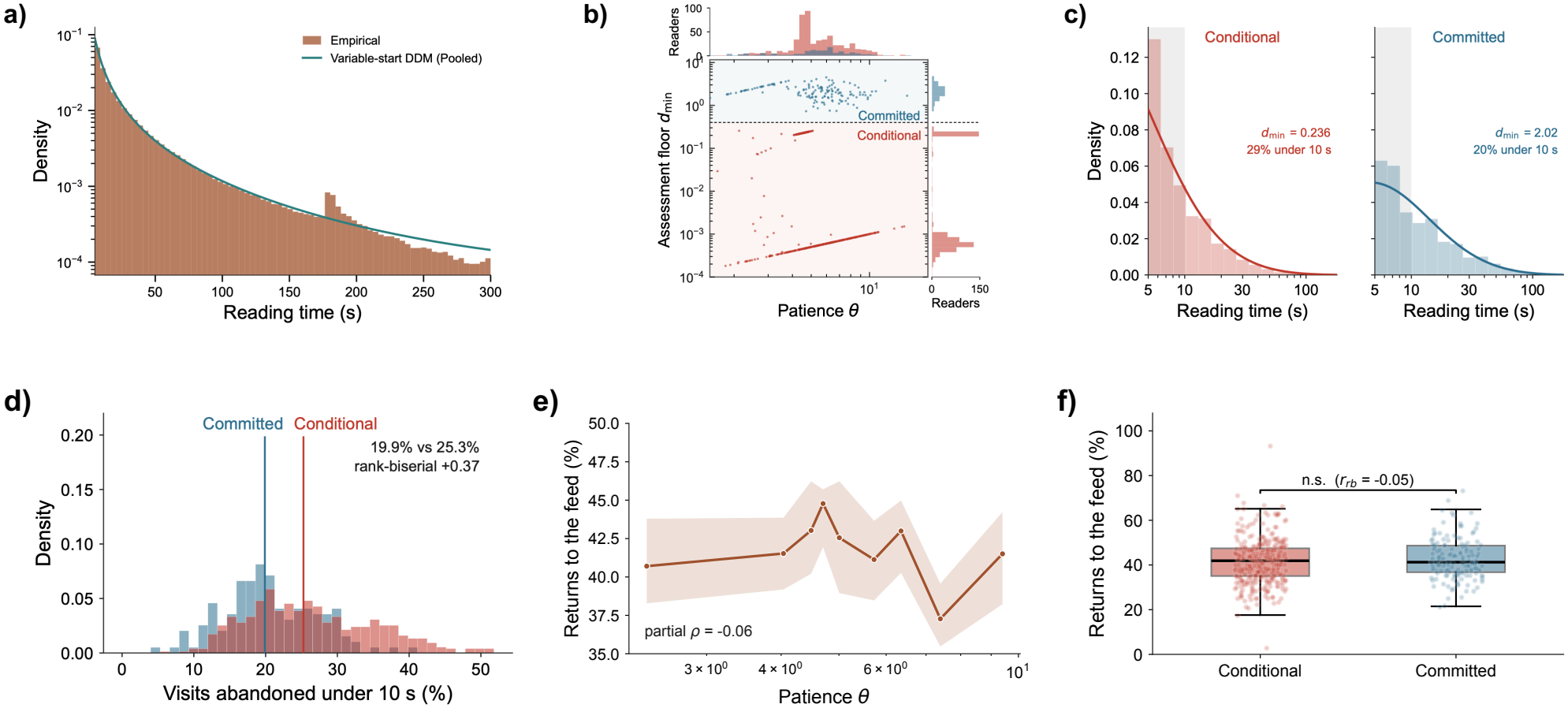
The assessment floor separates two groups of participants, with a weaker behavioural contrast in the new dataset. **(a)** Reading times pooled across all readers (green, *n* = 950678 reads), after excluding participants whose fitted start fraction reaches the lower bound, for whom the variable-start parameters are not interpretable, with the read-weighted average of the individually fitted densities, *p*(*t*) = ∑ _*i*_ *w*_*i*_*f*_*i*_(*t*), where *w*_*i*_ is proportional to the number of reads contributed by reader *i*. **(b)** Scatter plot of fitted parameters, assessment floor (*d*_min_) and patience (*θ*), of the variable-start model for each participant, with marginal distributions above and to the right. A Gaussian mixture on the fitted parameters, excluding the assessment floor itself, separates 717 participants into a group with an assessment floor close to zero (blue, *n* = 569, median *d*_min_ = 0.0007) and a group with a substantial assessment floor (red, *n* = 148, median *d*_min_ = 2.06). A third component of 67 participants, whose fits sit at the patience bound, is excluded as in the main analysis (Methods). The dashed line marks the boundary between groups. The assessment floor is bimodal across participants, whereas the two groups are not distinguished by patience. **(c)** Reading times of one representative participant from each group, conditional (red, left) and committed (blue, right), shown as histograms with their fitted densities. The two groups differ in the shape of their visit-time distributions in the way predicted by the assessment floor, with the difference concentrated in the shortest visits, as in the main analysis **(d)** Distribution across participants of the percentage of visits ending within 10 s, for the group with a substantial assessment floor (blue) and the group with an assessment floor close to zero (red). The groups differ in this share, 25.3% against 19.9%, respectively (rank-biserial correlation +0.37, *P* = 2 × 10^−12^). The direction is consistent with the main analysis, where the corresponding shares are 9.4% and 4.0%, respectively (rank-biserial correlation +0.77). **(e)** Median share of views classified as navigational, by quintile of patience. The shaded band represents the bootstrap 95% confidence interval of the median. The partial rank correlation between patience and navigational share, controlling for the assessment floor, is −0.06. **(e)** Share of views classified as navigational, for the group with a substantial assessment floor (blue) and the group with an assessment floor close to zero (red). The groups do not differ in this share, 41.3% against 41.9%, respectively (rank-biserial correlation −0.05, *P* = 0.35).

The behavioral patterns identified so far rest on a single population (Adressa dataset, Norway). To test whether they generalise to other kinds of digital content, we analysed an independent dataset, the Kulshrestha German web-tracking panel, composed of 2148 participants recorded over one month Kulshrestha et al. [2021], with 9.15 million page views across 49918 domains. The URLs in this dataset are anonymised, so pages cannot be classified by their address and navigational pages have to be identified from behaviour alone.

The Adressa analysis separates articles from front and section pages by their address, and no equivalent is available in this dataset. Here, 2148 participants are spread over 49918 domains, so most domains are visited by too few people for any rule that relies on many readers visiting the same page. We therefore adapted the maximal forward reference procedure of Chen, Park and Yu Chen et al. [1998], which segments a browsing session into forward paths and discards backward references on the grounds that these are made for ease of traveling rather than to consume content. The algorithm classifies no pages, so the extension here is to count, for each reader, how many of their own forward paths contain a given page. Counting within a reader rather than across readers is what makes the rule applicable here. A page’s classification depends on how that specific reader uses it, and the count is available whether a domain has two readers or several thousand.

Validated against Adressa, where the URL gives the answer, the maximal forward reference rule produces 96.9% balanced accuracy when pages appearing in fewer than three of a reader’s paths are treated as content, recovering 97.1% of articles and 96.7% of front pages. Cuts between two and ten all give between 92% and 97%, so the threshold is not carrying the classification. A dwell-time cutoff in the manner of the reference length method Cooley et al. [1999], with the assumed share of navigational references set from the Adressa labels themselves, produces 63.8% on the same test in comparison.Page views were processed as in the main analysis. Consecutive views of the same page (with the same url) within 30 s were merged into a single visit (5.6% of views, against 1.7% on Adressa), sessions were separated at a pause of 300 s, and the remaining visits were restricted to the [5, 300) s window. This leaves 1642585 visits from 2141 participants, of which 1265 contribute at least 300 visits and 968 at least 500. Applied here, the maximal forward reference rule classifies 32% of views as content against 35% on Adressa. However, it does not separate page durations, with both the retained views and the discarded ones having a median of 8 s, whereas on Adressa the discarded views are substantially the shorter.

Pooled across the retained visits, the distribution of visit times is described by accumulation from a variable starting point (Supplementary Fig. S12a), and the model returns a median patience of *θ* = 4.20 across the 1265 participants with at least 300 visits, against 17.35 on Adressa. The fitted density follows the empirical one with one systematic departure, a shoulder at 180 s, where the observed density rises above the fitted curve and then falls below it within a few seconds, a feature the model has no mechanism to produce. Moreover, engagement times in this dataset fall on even values far more often than chance, 82% against 49% in Adressa. A second difference from Adressa is in what is being visited. The Adressa dataset is composed of a single newspaper, so every retained visit is to an article. Here the retained visits span the whole of a participant’s browsing (news sites, video platforms, social networks, retailers, forums, web mail, etc.). A page on these is scrolled, watched or searched as readily as it is read, and the leaving decision the model describes need not govern how long a participant stays. The quantity fitted below is therefore the duration of a page visit rather than of an article read, and the parameters should be read with that in mind.

The start fraction is bimodal at the bounds, with 37% of participants reaching the upper bound, and a further 38% reaching the lower one, where the model reduces to a constant start and every visit requires the same evidence. On Adressa, 54% reach the upper bound and only 3% the lower. The participants reaching the lower bound are not badly fitted. Between each participant’s visits and visits simulated at their own fitted values, the Kolmogorov-Smirnov statistic has a median of 0.078 and no value above 0.15. Furthermore, twenty optimiser restarts return the same solution to four decimal places, and lowering the bound from 0.01 to 10^−4^ moves where the estimates accumulate without dispersing them. For these participants the data provide no evidence of evidence gathered before the visit, so the variable-start parameters are not interpretable and they are excluded, as participants at the opposite bound are treated separately in the main analysis. This removes more than a third of the sample, and the behavioural result reported below does not hold without it.

Across the 787 remaining participants the assessment floor is again bimodal. A Gaussian mixture on the fitted parameters, which do not include the floor itself, separates them into 569 with a floor close to zero (median *d*_min_ = 0.0007) and 148 with a substantial one (median 2.06). A third component of 67 participants, whose fits sit at the patience bound, is excluded as in the main analysis (Supplementary Fig. S12b). The group medians are similar to those obtained on Adressa, 0.0016 and 3.12. The two groups differ in the shape of their visit-time distributions in the way the assessment floor predicts, with the difference concentrated in the shortest visits (Supplementary Fig. S12c). They also differ in the share of visits ending within 10 s, 25.3% against 19.9%, a rank-biserial correlation of +0.37 (*P* = 2 × 10^−12^, Supplementary Fig. S12d). The direction is the one the main analysis reports, where the same contrast is 9.4% against 4.0% at a rank-biserial of +0.77. Computed over all 1265 participants, without the exclusion, the correlation is −0.10, a small effect in the opposite direction.

The navigational behaviour does not separate the groups. The analogue of returning to the front page is here inferred from the paths rather than read from the URL. That share slightly declines across the range of patience (partial rank correlation − 0.06 controlling for the assessment floor, Supplementary Fig. S12e), but the two groups do not differ on it, 41.9% against 41.3% (rank-biserial −0.05, *P* = 0.35, Supplementary Fig. S12f). On Adressa the corresponding partial correlation is −0.29.

We attribute the difference to what is being measured rather than to the participants. The quantity fitted here is the duration of a page visit on any site, not of an article read. News accounts for 2.6% of this panel’s browsing, and restricting the analysis to news domains leaves 8 participants above the threshold used in the main analysis, which is too few to fit. What survives the classification is dominated by social, retail and video sites, where a page is scrolled or watched rather than read, so the decision to leave is still being made, but over a different kind of content and plausibly on a different timescale. The median patience of 4.20 against 17.35, the failure of the classifier to separate page durations, and the absence of a navigational contrast are all consistent with this.

The division therefore recovers in a dataset collected in a different country, in a different year, by a different instrument, and over content that is largely not news, which argues that it is not an artefact of the Adressa dataset. Its behavioural signature is weaker and conditional on excluding participants for whom the start fraction is not identified. A dataset suitable for a full replication would require per-participant depth comparable to Adressa’s together with content that is predominantly articles.

## 7 Limitations of the study

Several limitations qualify the interpretation and generalisability of our results.

First, the main dataset comes from a single regional newspaper, in one language, over a ten-week period. Consequently, we cannot establish whether the behavioural division we identify reflects a general feature of readers’ information foraging behaviour or characteristics specific to this outlet and its audience. We examined the robustness of this finding using a second dataset collected in a different country, in a different year, using a different instrument, and covering content that was largely non-news. The persistence of a similar pattern suggests that our results are not simply an artifact of the Adressa dataset. However, the behavioural signature is weaker and conditional on excluding participants for whom the start fraction cannot be identified. A dataset suitable for a more comprehensive replication would need to combine per-participant observation depths comparable to those in Adressa with content consisting predominantly of news articles.

Second, the readers included in our analysis are, by construction, among the most active visitors to the site. Estimating three parameters for each reader requires sufficiently long behavioural records, which systematically excludes less engaged visitors. These readers may exhibit different foraging strategies, and their absence limits the extent to which our findings can be generalised to the newspaper’s entire readership. In particular, the observed behavioural division may partly reflect the characteristics of highly engaged readers rather than the full range of reading behaviours.

Third, the information-foraging framework distinguishes three related problems Pirolli and Card [1999]. The patch problem, concerning time allocation within information sources. The scent problem, concerning the evaluation of information cues, such as headlines, before accessing an article. The diet problem, concerning the selection of articles from the available alternatives. Our analysis directly models the patch problem by examining how readers allocate their attention within articles. We indirectly observe aspects of the scent problem through readers’ access patterns, but cannot fully decompose the processes underlying headline evaluation and article selection. We do not directly address the diet problem, since server logs do not provide a reliable reconstruction of the set of alternatives visible to each reader at any given moment. Consequently, our results characterise an important component of information foraging, rather than the complete decision-making process through which readers navigate an information environment.

Finally, the assessment floor is identified in terms of its ordering rather than its precise value for readers near the parameter bound. For these readers, the conditional group is therefore defined by the absence of a detectable assessment floor, rather than by an estimate indicating that the floor is small. This distinction matters for interpretation. The results establish a difference in the presence or detectability of an assessment floor, but do not necessarily imply that the underlying floor is negligible for every reader in this group.

